# Revealing long-range heterogeneous organization of nucleoproteins with N^6^-methyladenine footprinting

**DOI:** 10.1101/2024.12.05.627052

**Authors:** Wentao Yang, Xue Qing Wang, Aaron M Wenger, Fan Wei, Jingqi Yu, Yifan Liu, Yali Dou

## Abstract

A major challenge in epigenetics is uncovering the dynamic distribution of nucleosomes and other DNA-binding proteins, which plays a crucial role in regulating cellular functions. Established approaches such as ATAC-seq, ChIP-seq, and CUT&RUN provide valuable insights but are limited by the ensemble nature of their data, masking the cellular and molecular heterogeneity that is often functionally significant. Recently, long-read sequencing technologies, particularly Single Molecule, Real-Time (SMRT/PacBio) sequencing, have introduced transformative capabilities, such as N^6^-methyladenine (6mA) footprinting. This technique enables the detection of joint binding events and long-range chromatin organization on individual DNA molecules spanning multiple kilobases. Despite the potential of 6mA footprinting, existing analytical tools for 6mA detection in SMRT sequencing data suffer from significant limitations in both performance and applicability, especially with the latest sequencing platform. To address these gaps, we developed a novel 6mA-calling pipeline based on polymerase kinetics analysis. Our approach significantly outperforms current tools in terms of accuracy and computational efficiency, setting a new benchmark for 6mA detection. Utilizing our optimized experimental and computational framework, we extensively mapped nucleosome positioning and transcription factor occupancy at the single-molecule level, revealing critical features of the transcription-associated epigenetic landscape. Additionally, our work established high-resolution, long-range binding events in mitochondrial DNA, revealing simultaneous loading of two sets of replication machinery onto the displacement loop (D-loop). Our study highlights the potential of 6mA footprinting in capturing the coordinate binding of nucleoproteins and unraveling heterogeneous epigenetic states with unprecedented resolution.

## Introduction

One of the central tasks in epigenetics is to map the distribution of nucleosomes and other DNA-binding proteins across the genome [1, 2]. This is usually achieved by isolating DNA fragments corresponding to a specific nucleoprotein, which are then sequenced [3–6]. Various nucleases, such as DNase I, micrococcal nuclease (MNase), and Tn5 transposase, are routinely used for DNA fragmentation, either as a general probe of chromatin structure, in techniques like DNase-seq, MNase-seq, and ATAC-seq [7–9], or specifically targeted, as in CUT&RUN and CUT&Tag [10, 11]. The positional information of a nucleoprotein can be inferred from altered nuclease access, but the connectivity information between coordinate binding events on the same DNA molecule is lost. The derived data often need to be aggregated to reveal statistically significant patterns, which obscure dynamic or heterogenous protein-DNA interactions. The position for a nucleoprotein can also be registered by altered access to a DNA methyltransferase (MTase) and corresponding changes in DNA methylation (*N*^6^-methyl adenine: 6mA; 5-methyl cytosine: 5mC), implemented in techniques like DamID and NOMe-seq [12, 13]. DNA methylation footprinting, when coupled with long-read sequencing technologies, such as Single Molecule, Real-Time (SMRT/PacBio) sequencing and Oxford Nanopore (ONT) sequencing, has the potential to reveal joint binding events and long-range organization on individual chromatin fibers [13–17]. Particularly promising is the combination of 6mA footprinting (6mA-FP) and SMRT sequencing. SMRT sequencing can call 6mA (as well as 5mC) on individual DNA molecules multi-kb in length [18, 19]. Mammalian genomic DNA lacks 6mA [20–23], so essentially all 6mA sites are attributable to 6mA-FP. Combined with the ubiquitous presence of adenine sites and application of promiscuous 6mA MTases, 6mA-FP may detect DNA binding events at the single molecule level and with near-basepair resolution. Variably implemented as chromatin fiber sequencing (fiber-seq) [14], single-molecule adenine methylated oligonucleosome sequencing assay (SAMOSA) [16], and SAMOSA by tagmentation (SAMOSA-Tag) [24], 6mA-FP has been employed to study global chromatin organization, mitochondrial DNA compaction, and the epigenetic landscape. However, effective and reliable detection of base modifications, including 6mA, by long-read sequencing technologies remains a challenge and an area of active research [25].

In SMRT sequencing, 6mA is distinguished from the unmodified adenine by its perturbation to DNA polymerase kinetics—specifically increase in the time between nucleotide incorporation, referred to as the inter-pulse duration (IPD) (Fig. 1A) [19, 26]. 6mA mapping is usually achieved at the ensemble/genomic level, by combining different DNA molecules covering the same genomic position to overcome random fluctuations in IPD [27–31]. To generates accurate base calling and IPD information at the single molecule level, subread level data from multiple passes of the same DNA template is combined in Circular Consensus Sequencing (CCS; also known as PacBio HiFi Sequencing) (Fig. 1A) [18, 19, 23, 26]. Even though several bioinformatic pipelines have been developed for 6mA calling with SMRT CCS data [14, 16, 32, 33], various issues limit their widespread adoption, and as a consequence, effective implementation of 6mA-FP. We previously developed a pipeline that processes the subread level data from the PacBio Sequel systems and generates highly accurate 6mA calls at the single molecule level, but at a steep computational cost [32]. Furthermore, it is not applicable for the latest PacBio Revio platform, which no longer outputs the sequence and polymerase kinetics data at the subread level [34]. Fibertools, the latest open-source pipeline [33], makes inaccurate and incomplete 6mA calls. There is a demand for broadly applicable, accurate, and cost-effective 6mA calling at the single molecule level.

**Figure 1.**
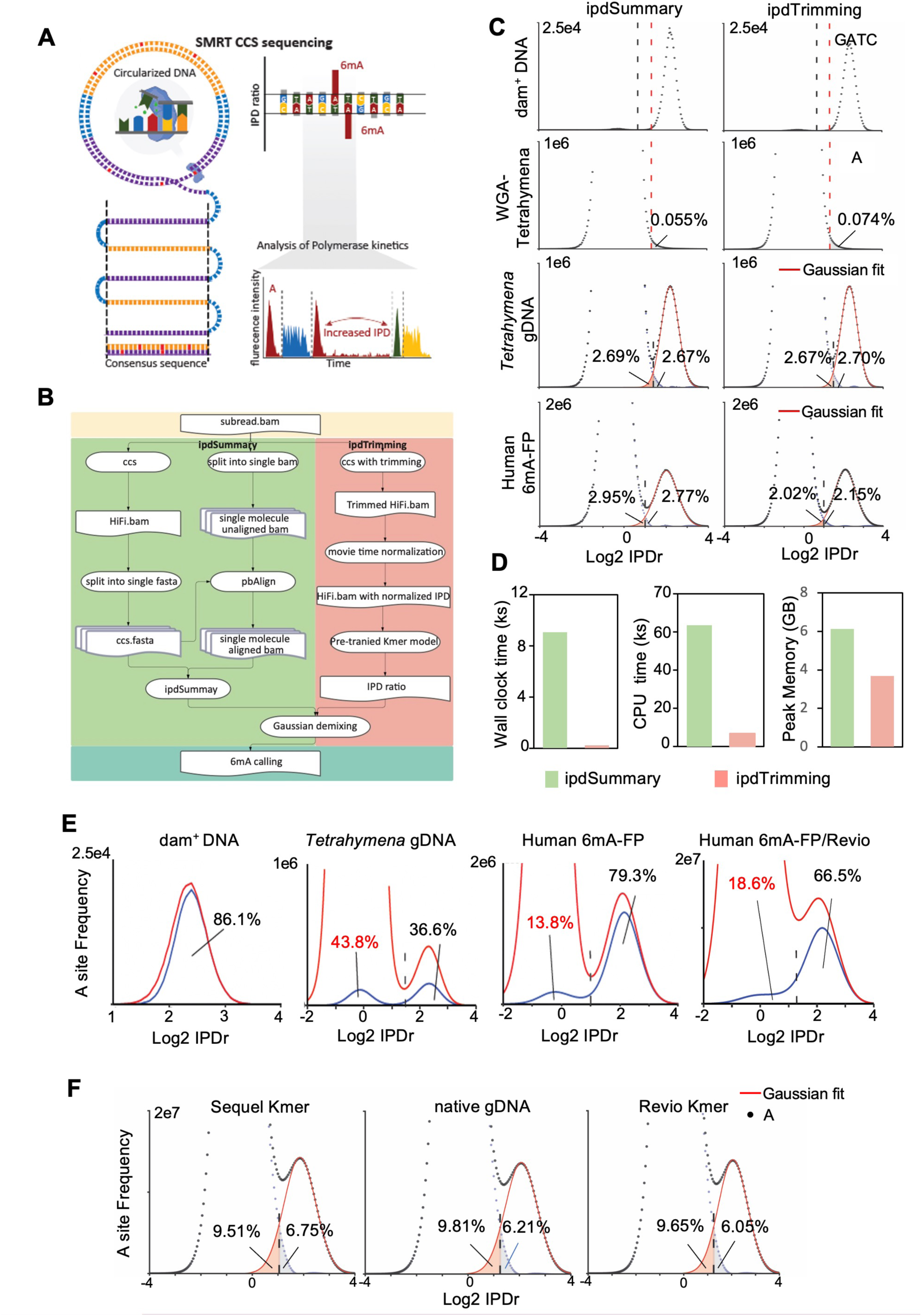
6mA calling for SMRT CCS. A. Schematic for SMRT CCS and 6mA calling. B. Comparing ipdSummary and ipdTrimming pipelines for 6mA calling using the subread level Sequel data. C. Performance of ipdSummary and ipdTrimming on four different datasets as indicated on left. The thresholds for calling 6mA are indicated by black lines. For dam^+^ plasmid DNA dataset, the thresholds for 98.5% 6mA recall are indicated by red lines. The same recall thresholds are used to calculate the background noise level for 6mA calling in the WGA dataset. False positive rates (FPR, grey shade) and false negative rates (FNR, red shade) for both *Tetrahymena* gDNA and human 6mA-FP samples were calculated after Gaussian demixing of IPDr distribution. D. Comparing the computational costs, including job all clock time, CPU running time and peak memory usage of ipdSummary and ipdTrimming. E. Comparing 6mA calling results by ipdTrimming and fibertools. Red curves represent IPDr distribution for all adenine sites calculated by ipdTrimming; blue curves represent the number of adenine sites with indicated IPDr values that are called as 6mA by fibertools. The thresholds for 6mA calling by ipdTrimming are indicated by black lines. Recall rates (black label) and FPR (red label) were calculated after Gaussian demixing of IPDr distribution. F. Comparing 6mA calling results for the Revio system, generated by the Sequel kmer model, human native gDNA control, and the Revio kmer model. FPR (grey shade) and FNR (red shade) were calculated after Gaussian demixing of IPDr distribution.

Here, we developed an innovative ipdTrimming pipeline that effectively controlled background noise in 6mA detection and achieved superior performance with the Sequel data at a minimal computational cost. Furthermore, this pipeline seamlessly adapted to the Revio platform. Critically, across a diverse range of samples sequenced on both Sequel and Revio systems, our method outperformed fibertools [33], reaching significantly higher precision and recall. Leveraging this cutting-edge bioinformatic workflow in conjunction with optimized 6mA-FP experimental procedures, we conducted detailed examinations of nucleoprotein organizations within human cells, encompassing both nuclear chromatin and mitochondrial nucleoid structures. Our analyses, at near-basepair resolution, revealed complex, long-range, and dynamic interactions involving nucleosomes, transcription factors/cofactors, and mitochondrial proteins. Our work positions 6mA-FP as a revolutionary tool for mapping the intricate organization of nucleoproteins and uncovering diverse epigenetic states at the single-molecule level.

## Results

### New 6mA calling pipelines for SMRT CCS

IPD conversion is a critical step in kinetics analysis of the Sequel data: for each site in a DNA molecule, multiple subread IPD values are converted to a single CCS IPD value (Fig. 1B). By default, subread IPD values are averaged to generate the CCS IPD value, routinely executed by the PacBio CCS module [34]. However, this default pipeline (referred to as CCS-kmer) generated inaccurate 6mA calls (Additional file 2: Table S1, S2). A major source of background noise in 6mA calling is DNA polymerase pausing, which stochastically introduces very large subread IPD values [35]. These outliers can substantially shift CCS IPD values, leading to false 6mA calls. The PacBio ipdSummary module, used for kinetics analysis at the ensemble level [34], controls this background noise by globally capping very large subread IPD values [35]. We have previously adapted ipdSummary for single molecule level 6mA calling, with high accuracy but at a steep computation cost [32]. Here, we developed a greatly simplified computation pipeline by introducing local trimming into IPD conversion: for each site in a DNA molecule, the top 10% of subread IPD values were removed before the rest were averaged to generate the CCS IPD value (Fig. 1B). The new IPD conversion step was executed by a modified CCS module. The CCS IPD value was subsequently compared to a reference IPD value of its unmodified counterpart embedded in the same local sequence; all reference IPD values were generated by a pretrained kmer model (https://github.com/PacificBiosciences/kineticsTools/tree/master/kineticsTools/resources). The derived IPD ratio (IPDr) was used for 6mA calling (Fig. 1B). The new pipeline, henceforth referred to as ipdTrimming, outperformed currently available 6mA calling methods.

We systematically compared CCS-kmer, ipdSummary, and ipdTrimming pipelines on several benchmarking datasets: ***1)*** a plasmid fragment from dam^+^ *E. coli*, with all A sites in GATC presumably methylated, serving as a positive control, ***2)*** *Tetrahymena* whole genome amplification (WGA), with all base modifications removed, serving as a negative control, ***3)*** native *Tetrahymena* genomic DNA, with <1% of all adenine sites methylated, ***4)*** 6mA-FP of in vitro assembled nucleosome arrays and human chromatin, with >10% of all adenine sites methylated (Fig. 1C, Additional file 1: Fig. S1, S2, and Additional file 2: Table S1). For the first two datasets, the IPDr for all A sites was distributed predominantly in a single peak, either high IPDr corresponding to 6mA or low IPDr corresponding to unmodified A (Fig. 1C, Additional file 1: Fig. S1). The background noise level for the WGA dataset was reduced from 0.37% for CCS-kmer to 0.074% for ipdTrimming, and close to 0.055% for ipdSummary (Fig. 1C, Additional file 1: S2). For the native *Tetrahymena* genomic DNA and 6mA-FP datasets that had both 6mA and unmodified A sites, IPDr exhibited a bimodal distribution: a large peak with low IPDr corresponding to unmodified A and a small peak with high IPDr corresponding to 6mA (Fig. 1C). We resolved the 6mA peak and unmodified A peak with Gaussian demixing and estimated the false positive and false negative rates (FPR and FNR) of 6mA calling methods (Fig. 1C, Additional file 2: Table S2). Both ipdSummary and ipdTrimming outperformed CCS-kmer; ipdTrimming performance was similar to ipdSummary at low 6mA levels (*Tetrahymena* dataset), but was substantially better at high 6mA levels (6mA-FP dataset).

We performed detailed comparison between ipdSummary and ipdTrimming using the native *Tetrahymena* genomic DNA dataset (Additional file 1: Fig. S2), as we have systematically characterized the endogenous 6mA sites by ipdSummary [32]. In all 6mA calling metrics, ipdTrimming closely matched ipdSummary (Fig. 1C, D, Additional file 1: Fig. S2, Additional file 2: Table S2). There was a very strong overlap between 6mA sites called by ipdTrimming and ipdSummary, with ipdTrimming calling slightly fewer 6mA sites (Additional file 1: Fig. S2). Furthermore, ipdTrimming recapitulated all key findings made with ipdSummary, including ***1)*** 6mA exclusively occurs at the ApT dinucleotide, ***2)*** full-6mApT is predominant over hemi-6mApT, and ***3)*** hemi-6mApT is predominantly found in only one strand of newly synthesized DNA duplex (Additional file 1: Fig. S2); all these results are consistent with semiconservative transmission of 6mA in this unicellular eukaryote [32].

To validate ipdTrimming for 6mA-FP, usually featuring much higher 6mA levels than natural occurrences, we first reanalyzed a previously published 6mA-FP dataset for an array of in vitro assembled nucleosomes (Additional file 1: Fig. S3, Additional file 2: Table S1) [16]. We found that the 6mA and unmodified A peaks were better resolved in ipdTrimming than ipdSummary (Additional file 1: Fig. S3, Additional file 2: Table S2). The vast majority of 6mA sites were jointly called by ipdTrimming and ipdSummary; they were depleted in nucleosome-protected regions (3.0% 6mA sites) and enriched in unprotected regions (97% 6mA sites) (Additional file 1: Fig. S3). Overall, 6mA calling by ipdTrimming was slightly more conservative than ipdSummary, and 6mA sites were slightly more depleted in nucleosome-protected regions (Additional file 1: Fig. S3). 6mA is known to have secondary offsite effects, most prominently at positions 5 nucleotides upstream (5’) of primary 6mA sites [36]. Intriguingly, ipdTrimming made substantially fewer 6mA calls at one of the likely secondary sites than ipdSummary (Additional file 1: Fig. S3). 6mA calls were further validated by cross-referencing structural and molecular dynamics results of the nucleosome (see below). We also examined datasets for 6mA-FP of human chromatin and found that the 6mA and unmodified A peaks were also better resolved in ipdTrimming than ipdSummary (Fig. 1C, Additional file 1: Fig. S3, Additional file 2: Table S2). The vast majority of 6mA sites were jointly called; all key features, including the ∼10 bp oscillation pattern, were preserved by both (see below) (Additional file 1: Fig. S3). Importantly, ipdTrimming had much lower computation cost than ipdSummary, in terms of job wall-clock time, CPU running time, and peak memory usage (0.023×, 0.11×, and 0.60×, respectively) (Fig. 1D). Therefore, ipdTrimming effectively supersedes ipdSummary for kinetic analysis of the Sequel data.

We also compared ipdTrimming with fibertools, the latest open-source pipeline developed for fiber-seq/6mA-FP [33]. With its default setting, fibertools only recalled 85.0% of GATC sites (presumably all 6mA) in the positive control dam^+^ plasmid DNA, as compared to 98.7% using ipdTrimming (Fig. 1E). Fibertools’ performance deteriorated rapidly with decreasing IPDr (Fig. 1E). For the native *Tetrahymena* genomic DNA, fibertools only recalled 36.6% of 6mA sites called by ipdTrimming, but falsely identified many unmodified A sites as 6mA (43.8% and 2.70% FPR for fibertools and ipdTrimming, respectively) (Fig. 1E). Close examination revealed that fibertools made 6mA calls at all four ApN dinucleotides, instead of exclusively at the ApT dinucleotide (Additional file 1: Fig. S4). For 6mA-FP of both in vitro assembled nucleosome arrays and human chromatin, fibertools provided a general profile consistent with the beads-on-a-string nucleosome distribution (Additional file 1: Fig. S4). Nonetheless, there were substantial discrepancies between 6mA calls made by ipdTrimming and fibertools (Fig. 1E, Additional file 1: Fig. S4). Importantly, fibertools exhibited some strong systematic biases: on the one hand, it tended to make false 6mA calls at sites proximal to true 6mA sites; on the other, it tended to miss isolated or sparsely deposited true 6mA sites (Additional file 1: Fig. S4). Analysis using fibertools led to loss/diminishing of many key features, including the ∼10 bp oscillation pattern signifying dynamics of nucleosomal DNA (Fig. 2E). We conclude that for the Sequel data, the ipdTrimming pipeline clearly outperforms fibertools in all metrics, establishing a new 6mA calling standard for both endogenous 6mA detection and 6mA-FP.

**Figure 2.**
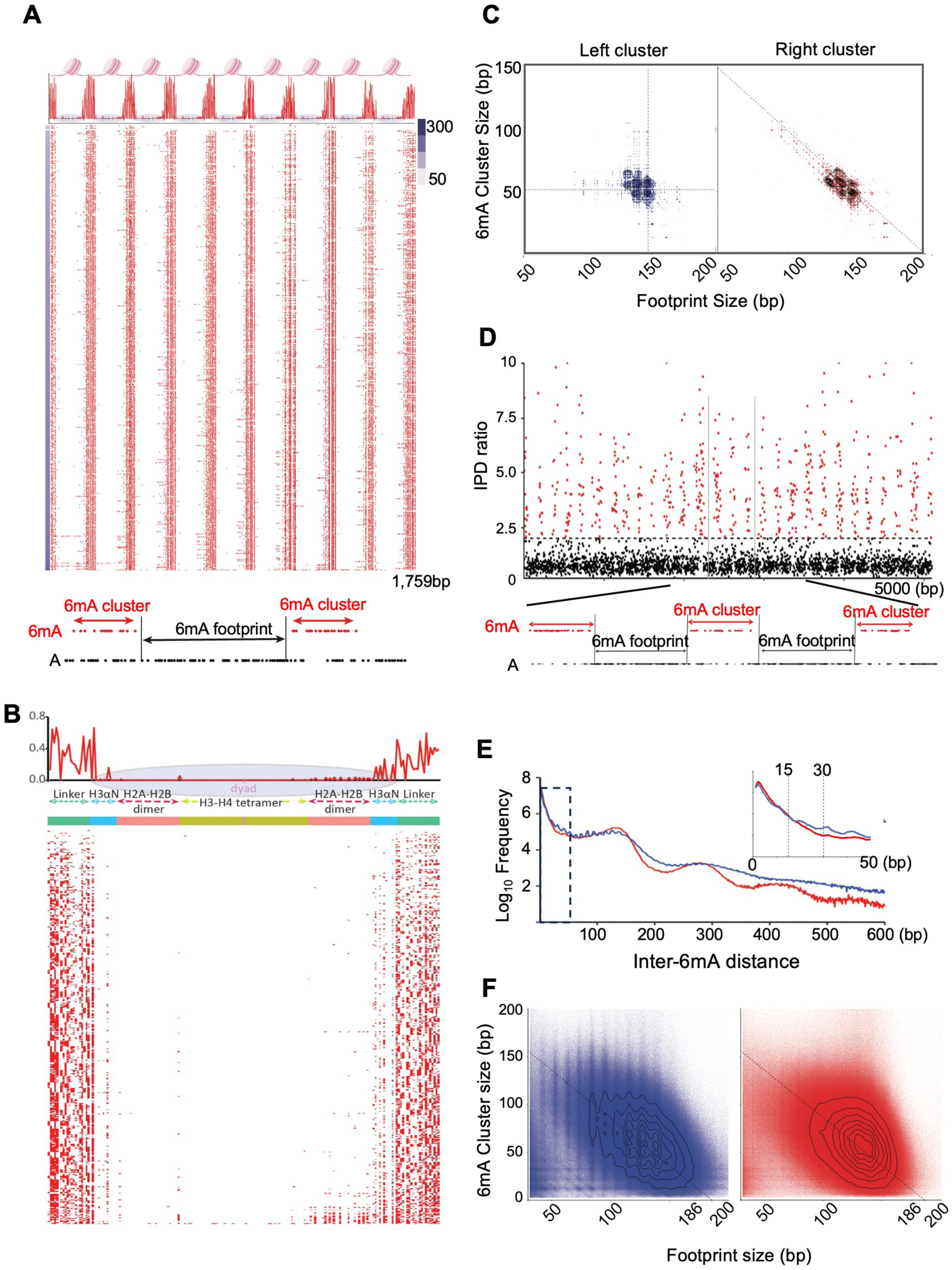
Nucleosome position and dynamics revealed by 6mA-FP. A. 6mA distribution on the Widom 601 nucleosome array after 6mA-FP. Top: a schematic diagram for positioned nucleosomes and linker DNA at regular intervals. Middle: 6mA distribution in aggregate (shown as 6mA penetrance) and in individual DNA molecules (sorted by the 6mA count). Bottom: zoom-in on a DNA molecule, illustrating the definition of the 6mA clusters and footprints. B. 6mA distribution on individual nucleosomes and their flanking linker DNA. Top: 6mA distribution in aggregate (shown as 6mA penetrance), superimposed on a schematic of a nucleosome positioned on the Widom 601 sequence; the nucleosome dyad, regions protected by H3-H4 tetramer, H2A-H2B dimer, H3 αN helix, as well as linker DNA are indicated. Bottom: 6mA sites on individual DNA molecules (sorted by the 6mA count). C. Relationship between the size of adjacent 6mA clusters and footprints. Note that for pairs with 6mA cluster on the left and nucleosome footprint on the right, horizontal and vertical stripes are prominent; for pairs with 6mA cluster on the right and nucleosome footprint on the left, diagonal stripes are prominent. D. Calling 6mA sites, 6mA clusters, and 6mA footprints on an individual human chromatin fiber after 6mA-FP. Top: IPDr values for all adenine sites on the DNA molecule. 6mA sites (red) and unmodified A sites (black) are separated by the IPDr threshold for 6mA calling (dashed line). Bottom: zoom-in illustrating the definition of the 6mA clusters and footprints. E. Distribution of the distance between adjacent 6mA sites on individual DNA molecules from short (blue) or long methylation (red) 6mA-FP of human chromatin. Most 6mA sites are close to each other (inset). Note major peaks in the long methylation (red) sample corresponding to the nucleosome ladder. F. Relationship between the size of adjacent 6mA clusters and footprints from 6mA-FP of human chromatin. Compare short (blue) and long (red) methylation. Note the inverse relationship between the size of adjacent 6mA clusters and footprints (dashed line) on both short (blue) and long (red) methylation; a grid-like pattern with ∼10 bp separation is only obvious for short methylation, corresponding to stepwise unwrapping of nucleosomal DNA.

For Revio, the latest PacBio platform for SMRT CCS, the on-instrument IPD conversion only outputs the CCS level IPD, by averaging all subread IPD values for each site of a DNA molecule. We explored using the Revio data for 6mA-FP (Fig. 1F, Additional File 1: Fig. S6). We first tested the kmer model pretrained on the Sequel data (Fig. 1F) (https://github.com/PacificBiosciences/kineticsTools/tree/master/kineticsTools/resources). The 6mA and unmodified A peaks for the Revio data were not as well separated as its Sequel counterparts, resulting in more erroneous calls (FPR: 6.75% and 1.67%, FNR: 9.51% and 1.68%, for Revio and Sequel, respectively) (Fig. 1C, F). To address the possibility that the performance issue was due to changes in chemistry, we generated a control Revio dataset of native human genomic DNA without 6mA-FP. We first assigned IPD values corresponding to the unmodified A for all relevant genomic positions, by averaging CCS IPD values of accurately mapped and aligned reads. We then compared IPD values of equivalent genomic positions from the 6mA-FP and control datasets to generate IPDr for 6mA calling. This approach (referred to as CCS-gDNA) yielded a moderate effect (FPR: 6.23%, FNR: 9.73%) (Fig. 1F). We then used the control Revio dataset to update the kmer model, by combining genomic positions with the same sequence context and averaging their IPD values. The new pipeline (referred to as CCS-kmer*) showed incremental improvement (FPR: 6.06%, FNR: 9.50%) (Fig. 1F). For comparison, fibertools had 66.5% recall rate and 18.6% FPR for the same Revio dataset (Fig. 1E). We conclude that our CCS-kmer* pipeline is the best available method for making 6mA calls and implementing 6mA-FP with the Revio system.

### Nucleosome position and dynamics revealed by 6mA-FP

As the basic unit of chromatin organization, the nucleosome features predominantly in 6mA-FP [14, 16]. To gain new insights into nucleosome positioning and dynamics, we used our ipdTrimming pipeline to analyze a previously published 6mA-FP dataset for an in vitro assembled nucleosome array (Fig. 2A, Additional file 1: Fig. S3) [16]. Linker DNA regions were densely methylated on individual DNA molecules, with many sites near saturation at the ensemble level (Fig. 2A, Additional file 1: Fig. S3). At the single molecule level, we aggregated multiple closely spaced 6mA sites into a 6mA cluster; complementarily, the interval between adjacent 6mA clusters, if containing multiple unmodified A sites, was designated a 6mA footprint (Fig 2A, bottom). 6mA clusters readily expanded beyond the linker DNA boundary and into the 147 bp Widom 601 sequence constituting the nucleosome core particle (Fig. 2A-C) [37]. Most expansions stayed in the terminal 13 bp of the Widom 601 sequence and stopped outside the central 121 bp nucleosomal DNA (nucleosome dyad ± 60 bp), supporting an octasome footprint with unwrapping of DNA in contact with the H3 αN helix and the root of the H3 N-terminal tail [38, 39]. Some 6mA clusters encroached into the region ±30 to ±60 bp from the nucleosome dyad, supporting an octasome-to-hexasome transition with unwrapping of DNA in contact with the H2A-H2B dimer [39]. Intriguingly, we also found a single A site on the left half (-31 bp from the dyad) with usually high 6mA penetrance (Fig. 2B), supporting dynamic opening of the interface between the H2A-H2B dimer and the H3-H4 tetramer [40]. Very rarely did they reach within ±30 bp from the nucleosome dyad, representing complete loss of H2A-H2B protection from either side or both. Notably, DNA unwrapping occurred preferentially from the right half of the Widom 601 nucleosome (Fig. 2B, C), consistent with tighter histone binding on the left half due to the presence of highly flexible pyrimidine-purine dinucleotides at the inward-facing minor groove regions [41, 42]. The disparity in unwrapping behaviors also manifested as two distinct relationships between a pair of adjacent 6mA cluster and 6mA footprint (Fig. 2C). For a 6mA cluster on the left of a Widom 601 nucleosome, its size was largely independent from the size of the 6mA footprint for the nucleosome, indicating that the left 6mA cluster rarely encroaches into the nucleosome on its right. Their size distribution plot featured prominent horizontal and vertical stripes intersecting at the most probable states, roughly corresponding to sizes of the linker and nucleosomal DNA (Fig. 2C). For a 6mA cluster on the right of a Widom 601 nucleosome, its size mostly covaried in a negative linear relationship with the size of the 6mA footprint for the nucleosome (Fig. 2C). The diagonal stripes, also passing through the most probable states, indicate that the right 6mA cluster frequently encroaches into the nucleosome on its left. Thus, by analyzing both 6mA clusters and the complementary 6mA footprints, we can accurately determine nucleosome position and dynamics in the Widom 601 nucleosome array.

To apply 6mA-FP to study human chromatin, we used M.EcoGII, a prokaryotic MTase targeting adenine sites in any sequence context [43], which was able to deposit 6mA at higher density and generate more prominent footprints with less sequence bias than Hia5 (Additional file 1: Fig. S7) [14, 33, 44]. Nuclei isolated from human acute myeloid leukemia (AML) cell lines MOLM13 and OCI-AML3 were subjected to short (1 hour) or long in vitro methylation (3 hours) by M.EcoGII (Additional file 1: Fig. S7) [45, 46]. After in vitro methylation, genomic DNA was digested by DpnI (targeting fully-methylated GATC sites) to generate fragments enriched for 6mA and of the preferred size range (3-5 kb) for accurate 6mA calling (Additional file 1: Fig. S7). The 6mA-FP samples were sequenced on both Sequel and Revio systems, and 6mA sites were called by ipdTrimming and CCS-kmer*, respectively. For 6mA-FP of human chromatin, 6mA sites on individual DNA molecules showed a very strong tendency of clustering, and most adjacent 6mA sites were separated by a distance smaller than the linker DNA length (78.8% ≤15bp, 84.1% ≤25bp, 86.8% ≤35bp) (Fig. 2D, E). 6mA clusters correspond to accessible DNA, with small ones representing linker DNA and large ones representing nucleosome-depleted regions (NDR) (Fig. 2D). Complementarily, 6mA footprints correspond to inaccessible DNA, with large ones protected by the nucleosome and small ones protected by other chromatin-associated proteins, including TFs and RNA polymerases (Fig. 2D). We plotted the size distribution of all pairs of adjacent 6mA cluster and footprint in short or long methylation samples (Fig. 2F). For short methylation 6mA-FP, the distribution showed a mixed pattern: ***1)*** a diagonal feature was generated by 6mA clusters encroaching into nucleosomal 6mA footprints, ***2)*** horizontal and vertical stripes were indicative of resistance to unwrapping of nucleosomal DNA, with a 10-bp grid consistent with stepwise unwrapping (Fig. 2F). For long methylation 6mA-FP, the diagonal feature became more prominent but also more diffusive, while the horizontal/vertical feature was diminished, as resistance to unwrapping was overcome by nucleosome dynamics over time (Fig. 2F). These results are consistent with the model based on the Widom 601 nucleosome array (Fig. 2C), and validate that 6mA-FP can effectively reveal nucleosome position and dynamics in human chromatin.

### Regularity and period of nucleosome arrangements in human cells revealed by 6mA-FP

Autocorrelation analysis of 6mA sites showed periodic distribution on most DNA molecules, with large oscillating periods of ∼200 bp (Fig. 3A). This pattern is underpinned by the beads-on-a-string arrangement of nucleosomes constituting the 10-nm chromatin fiber, with the period for 6mA oscillation corresponding to the nucleosome repeat length (NRL) [47]. Nonetheless, 6mA oscillation exhibited substantial variations in its regularity and period among individual DNA molecules (Fig. 3A). We performed spectral clustering of auto-correlograms to distinguish five classes of DNA molecules (Fig. 3A-C) [48]. Interestingly, Class I molecules showed low regularity and broad period in 6mA distribution (Fig. 3C). In contrast, Class II-V molecules had a more regular 6mA distribution and progressively longer 6mA period, likely corresponding to NRL of approximately 179, 189, 195, and 204 bp, respectively (Fig. 3C). Indeed, regularity and period were the two principal determinants for classification based on spectral clustering (Fig. 3D). Functional annotation showed that Class I molecules were enriched for promoters and enhancers of active genes, but depleted for repetitive sequences (Fig. 3E, I, Fig. 6F). This is consistent with chromatin opening and nucleosome disruption at transcription regulatory elements, as opposed to the more closed heterochromatin regions with more regular nucleosome arrays [49].

**Figure 3.**
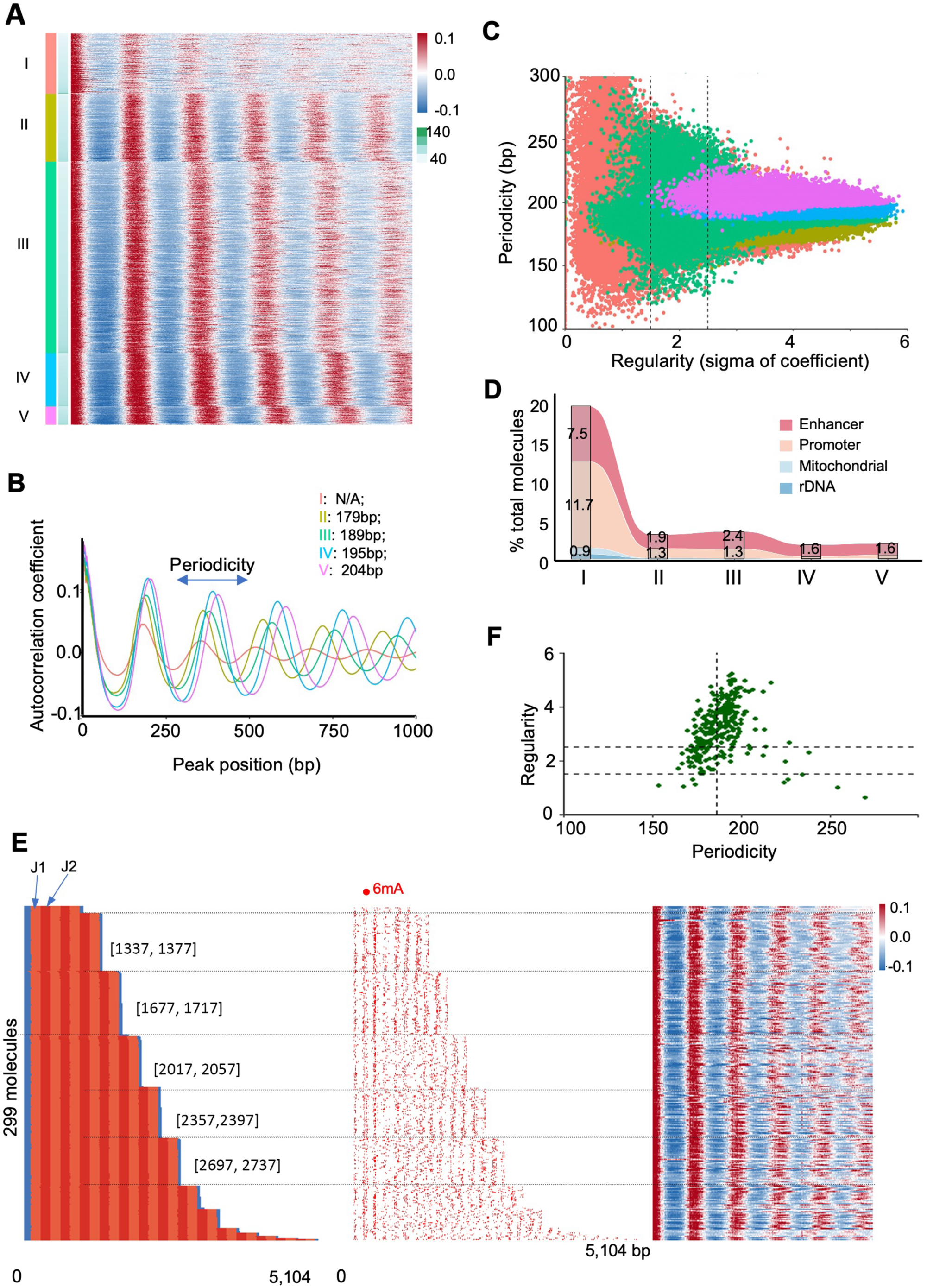
Heterogeneity in human chromatin revealed by 6mA-FP. A. Autocorrelograms of 10,000 randomly sampled DNA molecules, classified by spectral clustering (Class I-V) and ranked by 6mA counts. Top color bar: autocorrelation coefficient σ; bottom color bar: 6mA density (count/kb). B. Aggregated correlograms for each class. Nucleosome repeat length (NRL) estimation is shown for cluster II-V. C. Scatter plot showing the relationship between regularity (sigma of autocorrelation coefficient σ) and period (estimated NRLs) at the single-molecule level (color-coded for each cluster). D. Alluvial plot showing the proportional shifts of annotated molecules across each cluster. E. 6mA signals at J1/J2 centromeric higher-order repeat (HOR) J1/J2. DNA molecules of J1/J2 HOR (color-coded for J1/J2 monomer, left), 6mA sites on individual DNA molecules (middle), and autocorrelation between 6mA sites (right) from 6mA-FP of human chromatin, ranked by read length. F. Scatter plot showing the relationship between 6mA period (x-axis) and regularity (y-axis) in J1/J2 reads. Vertical dash line indicates 186bp of average NRL.

It has long been assumed that α-satellite repeats constituting the centromeric heterochromatin can strongly position nucleosomes [50]. 6mA-FP allowed us to examine chromatin organization at human centromere, fully sequenced only recently [51], at the single molecule level. Most centromeric molecules had regular nucleosome distribution, corresponding to Class II-V as specified above (Fig 3F). Unexpectedly, we found substantial heterogeneity in 6mA periods among DNA molecules mapped to the representative α-satellite higher-order repeat (HOR) J1/J2, despite essentially the same underlying sequence (Fig. 3F-I, Additional file 1: Fig. S8). This was reflected by the diversity in their classification by spectral clustering (Fig. 3I). Furthermore, nucleosome positioning was only observed near the ends of these DNA molecules, but it decayed very quickly inside (Fig. 3G). This result illustrates 6mA-FP’s power to reveal the intrinsic heterogeneity in chromatin organization at the single molecule level, which is often underestimated if only extrapolating from the ensemble level data.

### Transcription factor (TF)-binding events and transcription-associated epigenetic landscape

Combinatorial binding of transcription factors (TFs) at promoters and enhancers controls transcription in eukaryotes [52–54]. TF usually resides in a nucleosome-depleted region (NDR) [54]. Its binding can lead to MTase protection, leaving behind a small footprint nested inside the NDR (Fig. 4A: left). However, TF binding is often very dynamic, and its MTase protection is readily overcome during 6mA-FP, leaving behind only a large 6mA cluster encompassing the NDR (Fig. 4A: right). To examine whether our optimized 6mA-FP procedure can detect dynamic TF binding events at the single molecule level, we first focused on CTCF, for the following reasons: ***1)*** CTCF recognizes a long sequence motif and has potentially a larger footprint than most TFs [55–59]; ***2)*** stably bound CTCF, likely in association with the cohesin complex [60], has much longer residence time (in minutes) than most TFs (in seconds) [59, 61, 62]; and ***3)*** high-quality ChIP-seq data for CTCF are available in the human cell lines we studied (MOLM13 and OCI-AML3) [63–65]. To distinguish strong and weak CTCF binding sites, we ranked all genomic positions with a high-confidence CTCF binding motif by their ChIP-seq signals (Fig. 4B, E). DNA molecules covering strong CTCF binding sites generally featured a small 6mA footprint, <50 bp in size, centered around the CTCF consensus sequence. These CTCF footprints were 5mC-hypomethylated, demarcated by densely methylated 6mA clusters, and often flanked by large 6mA footprints corresponding to well-positioned nucleosomes (Fig. 4B-D, Additional file 1: Fig. S9). In contrast, DNA molecules covering weak CTCF binding sites showed few small 6mA footprints centered around the CTCF consensus sequence, which were generally 5mC-hypermethylated (Fig. 4B-D, Additional file 1: Fig. S9). Moreover, nucleosome-sized 6mA footprints were randomly distributed across these weak CTCF binding sites, without any strong positioning (Fig. 4B-D, Additional file 1: Fig. S9). Prolonged treatment by M.EcoGII largely eliminated CTCF footprints at the strong binding sites and merged adjacent 6mA clusters into a large one, while few large 6mA clusters were detected in DNA molecules covering the weak binding sites (Fig. 4E-G, Additional file 1: Fig. S9). Indeed, DNA molecules with strong or weak CTCF binding exhibited clear visual contrast both individually and in aggregate (Fig. 4B-G). We conclude that 6mA-FP readily reveals the dynamic CTCF binding events at the single molecule level, as either large 6mA clusters or small 6mA footprints, often accompanied by phasing of flanking nucleosomes.

**Figure 4.**
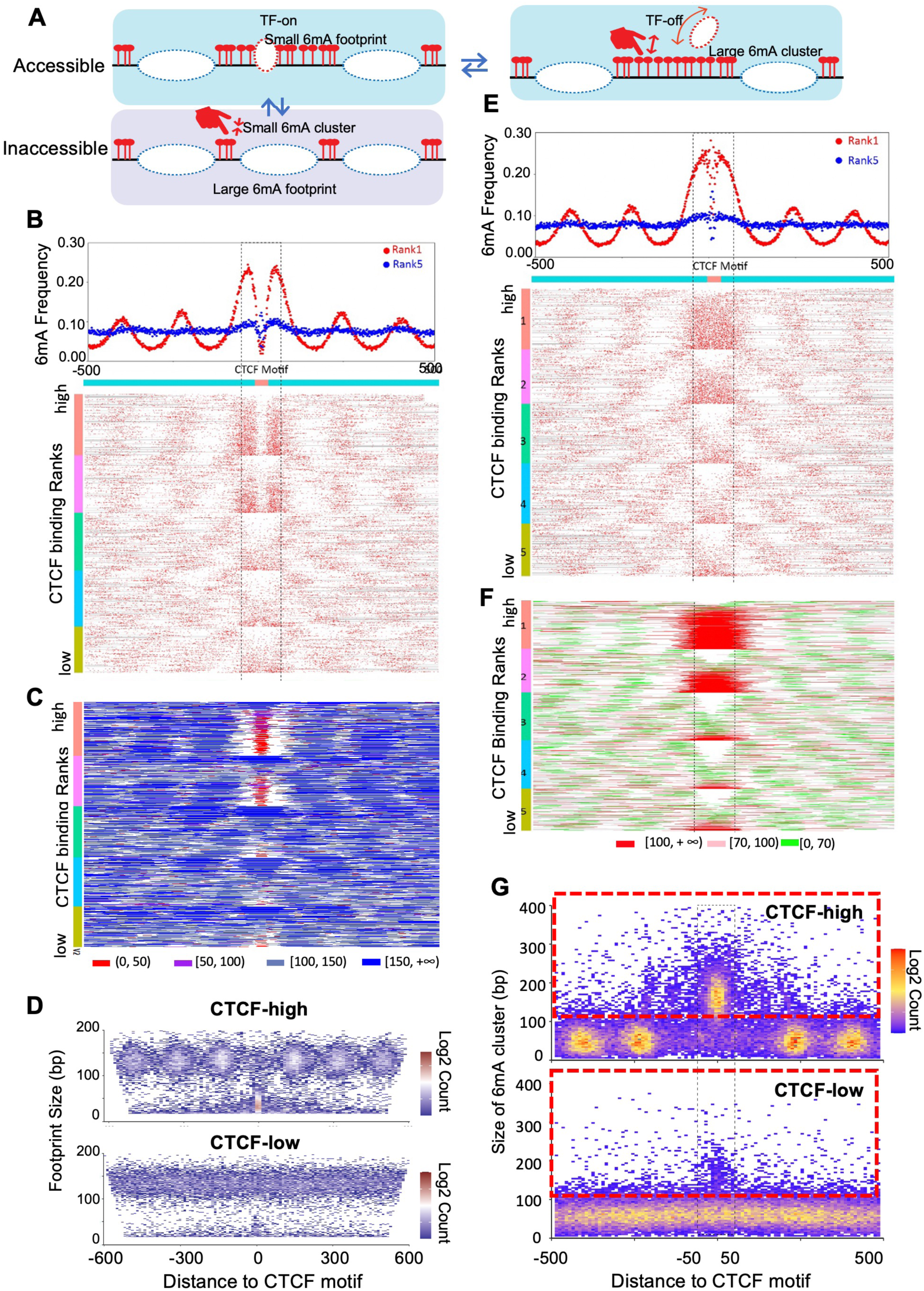
CTCF-binding events revealed by 6mA-FP. A. Schematic for detecting TF binding by 6mA-FP. Small 6mA clusters indicate linker DNA and chromatin regions inaccessible to TFs; small 6mA footprints nested inside NDR indicate the TF-on state of chromatin regions accessible to TFs; large 6mA clusters encompassing NDR indicate the TF-off state of chromatin regions accessible to TFs. B. CTCF binding events revealed by short methylation 6mA-FP. 6mA sites on individual DNA molecules centered around the CTCF binding motif are shown on the bottom. They were ranked from highest to lowest by CTCF ChIP-seq read counts (for a 118 bp window centered at the CTCF binding motif). Aggregated 6mA distribution around the CTCF binding motif are shown for Rank 1 and Rank 5 DNA molecules on top. C. Small 6mA footprints revealed by short methylation 6mA-FP. DNA molecules are ranked as in Fig. 4B. 6mA footprints are color coded by the size, shown at bottom. D. Heatmap for distribution of size (y-axis) and position (x-axis, relative to the CTCF binding motif) of 6mA footprints in CTCF-high (Rank 1) and CTCF-low (Rank 5) DNA molecules. Most footprints are between 100 to 200 bp, corresponding to nucleosomes. Note the strong enrichment of small 6mA footprints (≤50 bp), corresponding to the CTCF footprints, around the center in CTCF-high but not CTCF-low DNA molecules. E. CTCF binding events revealed by long methylation 6mA-FP. Treated the same as Fig. 4B. F. Large 6mA clusters revealed by long methylation 6mA-FP. DNA molecules are ranked as in Fig. 4E. 6mA clusters are color coded by the size, shown at bottom. G. Heatmap for distribution of size (y-axis) and position (x-axis, relative to the CTCF binding motif) of 6mA clusters in CTCF-high (Rank 1) and CTCF-low (Rank 5) DNA molecules. Most 6mA clusters are between 30 to 100 bp, corresponding to linker DNA. Note the strong enrichment of large 6mA clusters (≥100 bp), corresponding to NDR, around the center in CTCF-high but not CTCF-low DNA molecules.

We extended our analyses to other transcription factors (TFs) highly expressed and functionally important in the human AML cell lines OCI-AML3 and MOLM13, including SPI1/PU.1, CEBPα, MEF2C, MEF2D, IRF8, and MEIS1. By examining their ChIP signals in the two cell lines, we found that these TFs were mostly enriched in and around large 6mA clusters (≥100 bp) that were rare, but not small 6mA clusters (50-60 bp) that were more prevalent (Fig. 5A, B, Additional file 1: Fig. S10). This probably reflects the much shorter chromatin residence time for most TFs, relative to the CTCF residence time as well as the in vitro methylation time for 6mA-FP. Therefore, large 6mA clusters, rather than small 6mA footprints, are a much more sensitive, reliable, and quantitative binding indicator for most TFs. Furthermore, only large 6mA clusters were preferentially flanked by active histone marks associated with promoters and enhancers, including histone H3K4 methylation and multiple histone acetylation events (Additional file 1: Fig. S10). In contrast, no preference for repressive histone marks was found (Additional file 1: Fig. S10).

**Figure 5.**
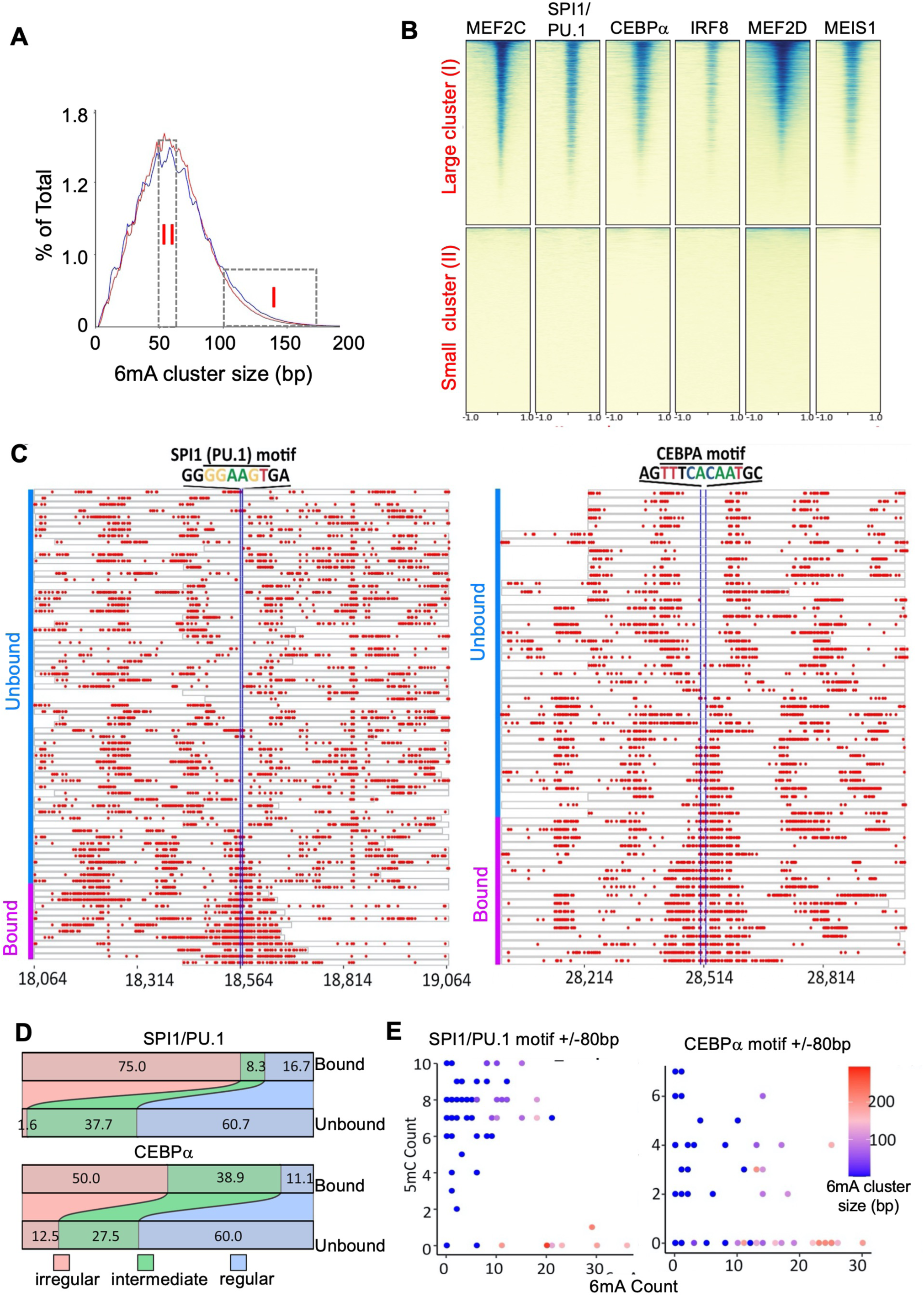
TF binding at large 6mA clusters. A. Size distribution of 6mA clusters. Two classes of 6mA clusters are defined according to their sizes: Class I (≥100 bp) and Class II (60 bp ≥ size ≥50 bp). B. Heatmap illustrating differential binding (ChIP-seq data for multiple TFs) in large (I) and small 6mA clusters (II). C. SPI1/PU.1 (left) and CEBPα (right) binding detected at single locus in human rDNA. DNA molecules spanning SPI1/PU.1 or CEBPα binding motif are sorted by the size of the 6mA cluster overlapping with the motif. The TF binding states are indicated on the left. D. Alluvial plot showing the shift in 6mA regularity for DNA molecules in the bound or unbound state. E. Scatter plot showing the inverse relationship between 6mA and 5mC count around SPI1/PU.1 and CEBPα binding motif. Color bar indicates the 6mA cluster size overlapping with the motif.

6mA-FP has the potential to fully reveal heterogeneity in TF binding at the single molecule level. To this end, we examined SPI1/PU.1 and CEBPα binding on 28S rDNA, which was deeply covered by SMRT CCS due to its high copy number (Fig. 5C). Strong SPI1/PU.1 and CEBPα binding in the intergenic spacer region of 28S rDNA has been reported in the leukemia cells, with important roles in modulating nucleolar transcription and cell proliferation [66]. Here we found that for DNA molecules covering the same genomic locus, only a fraction of them contained an overlapping large 6mA cluster, which disrupted the regular 6mA pattern associated with nucleosome arrays (Fig. 5C-E, Additional file 1: Fig. S11). In contrast, the rest of the DNA molecules contained no (large) 6mA clusters at the binding motif, showed predominantly 6mA patterns indicative of regularly spaced—but randomly positioned—nucleosomes (Fig. 5C-E). An inverse relationship between exogenous 6mA and endogenous 5mC was also detected for both SPI1/PU.1 and CEBPα binding loci at the single molecule level (Fig. 5D, Additional file 1: Fig. S11). Our result is consistent with SPI1/PU.1 and CEBPα binding at selective rDNA copies and subsequent recruitment of ATP-dependent chromatin remodeling and 5mC demethylation activities [67–70], even though we cannot completely rule out the possibility that they directly bind nucleosomal DNA and minimally disturb the local 6mA pattern. These results demonstrate 6mA-FP’s power to dissect the heterogeneity in TF binding and related events leading to transcription activation.

Regulatory signals for RNA polymerase II (Pol II) transcription are integrated through recruitment of transcription factors/cofactors, chromatin modifying/remodeling activities, as well as transcriptional/co-transcriptional machineries, generating specific transcriptional output in coordination with the epigenetic landscape [71]. We closely examined 6mA-FP data related to transcription-associated epigenetic changes. We focused on transcriptional start sites (TSS) of Pol II-transcribed genes, which were ranked and classified by their epigenetic or transcriptional states (Fig. 6A, D). 6mA-FP revealed unusual patterns in 6mA clusters and footprints associated with genes enriched for two active histone marks—H3K4me3 and H3K27ac (Fig. 6A). While there was significant heterogeneity, a large fraction of DNA molecules contained large 6mA clusters and/or small 6mA footprints around TSS, which was obvious at both the single molecule level and in aggregation (Fig. 6B, C). Regularity of nucleosome arrangements was also disrupted on DNA molecules mapped to these TSS, which predominantly belonged to Class I according to spectral clustering (Additional file 1: Fig. S12). In contrast, a much smaller fraction of DNA molecules exhibited a similar 6mA pattern for TSS that had either no H3K27ac or depleted for both marks, and showed irregular 6mA distribution (Fig. 6A-C). We also ranked TSS by their transcript levels and found that a similar bias for large 6mA clusters and/or small 6mA footprints to be associated with TSS of highly expressed genes (Fig. 6D-F). These large 6mA clusters, hypomethylated at 5mC, were attributed to NDR generated by chromatin opening at core/proximal promoters (Additional file 1: Fig. S12). While binding of TFs, including CTCF, certainly contributes to these small 6mA footprints, a more likely scenario is stably bound Pol II, especially the preinitiation complex (PIC) and promoter proximal pausing complex (PPP), as recently reported in a fiber-seq analysis of transcription in *Drosophila* [72].

**Figure 6.**
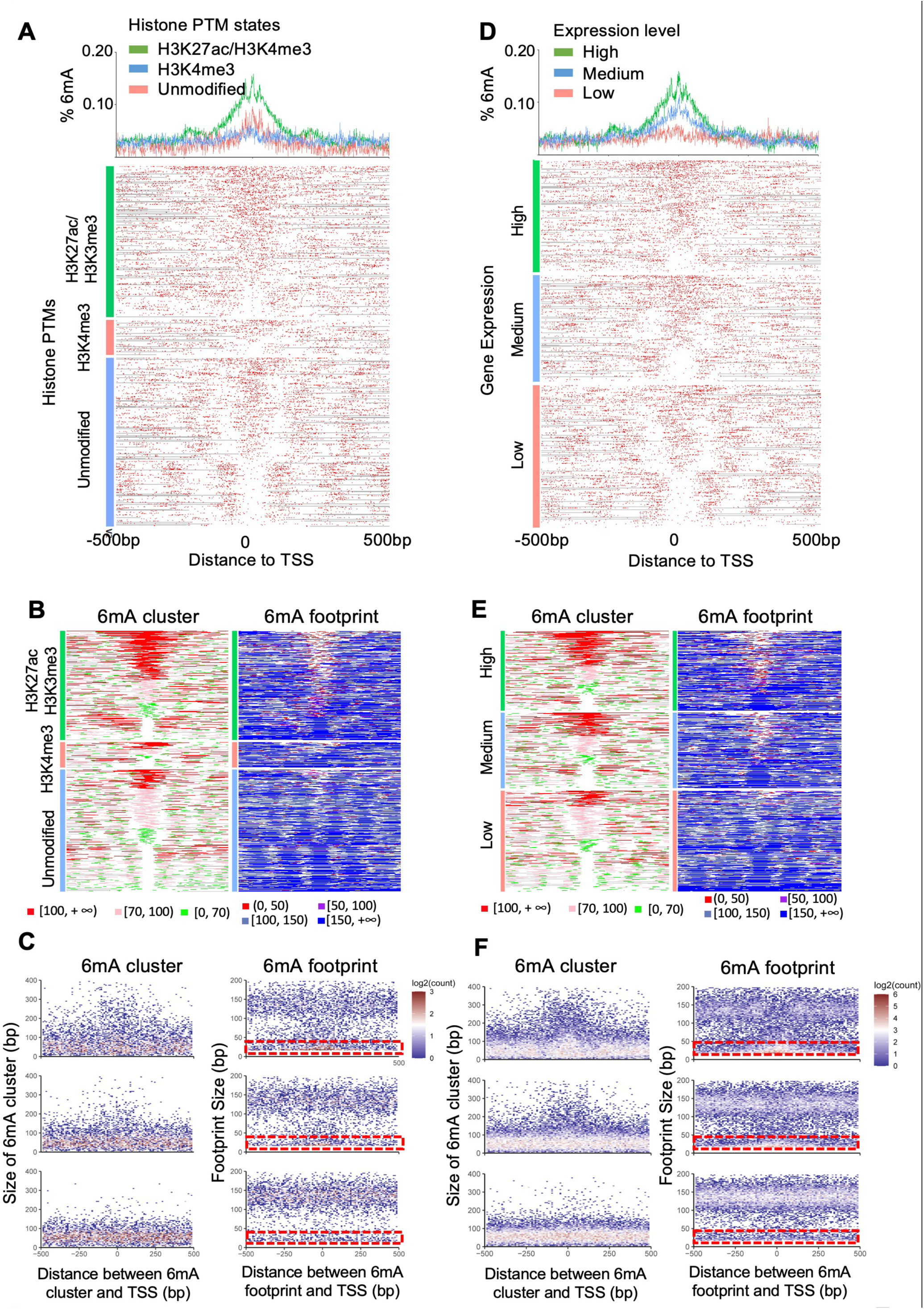
Altered chromatin organization around transcription start sites (TSS). A. 6mA distribution around TSS enriched for both H3K27ac and H3K4me3, only H3K4me3, or neither. Top: aggregated 6mA signals, centered around the three categories of TSS. Bottom: 6mA sites on individual DNA molecules, classified by their epigenetic marks and sorted by the size of the 6mA cluster overlapping with TSS. B. Distribution of 6mA clusters (left) and footprints (right) on individual molecules, classified and sorted as in Fig. 6A. Both 6mA clusters and footprints are color coded by their sizes. C. Heatmap for distribution of size (y-axis) and position (x-axis, relative to TSS) of 6mA clusters (left) and footprints (right) in DNA molecules for each category. Dashed boxes highlight small footprints (≤ 50 bp). D. 6mA distribution around TSS of genes with high, medium, or low expression levels. Top: aggregated 6mA signals, centered around the three categories of TSS. Bottom: 6mA sites on individual DNA molecules, classified by their expression levels and sorted by the size of the 6mA cluster overlapping with TSS. E. Distribution of 6mA clusters (left) and footprints (right) on individual molecules, classified and sorted as in Fig. 6D. Both 6mA clusters and footprints are color coded by their sizes. F. Heatmap for distribution of size (y-axis) and position (x-axis, relative to TSS) of 6mA clusters (left) and footprints (right) in DNA molecules for each category. Dashed boxes highlight small footprints (≤ 50 bp).

### Nucleoid organization of mitochondrial DNA (mtDNA)

The small circular mitochondrial DNA (mtDNA: 16.5 kb; Fig. 7A) is present at hundreds to thousands of copies in human cells [73]. Nuclear-embedded mitochondrial DNA (NUMT), some resulting from recent and ongoing transfer of mtDNA into the nucleus, complicates the analysis of heteroplasmic mtDNA [74, 75]. Here we show that 6mA-FP can effectively distinguish NUMT and mtDNA by detecting their unique genetic and epigenetic signatures (Fig. 7B). Consistent with its nucleosome-free nature, mtDNA was densely decorated with 6mA, with very few nucleosome-sized 6mA footprints (Fig. 7C). In contrast, NUMT segments featured the periodic 6mA pattern indicative of the nucleosome-based chromatin organization (Fig. 7C). NUMT segments were further distinguished by high number of 5mC calls and mismatches, consistent with their nuclear presence and high mutation rates due to lack of selection (Fig. 7C, inset).

**Figure 7.**
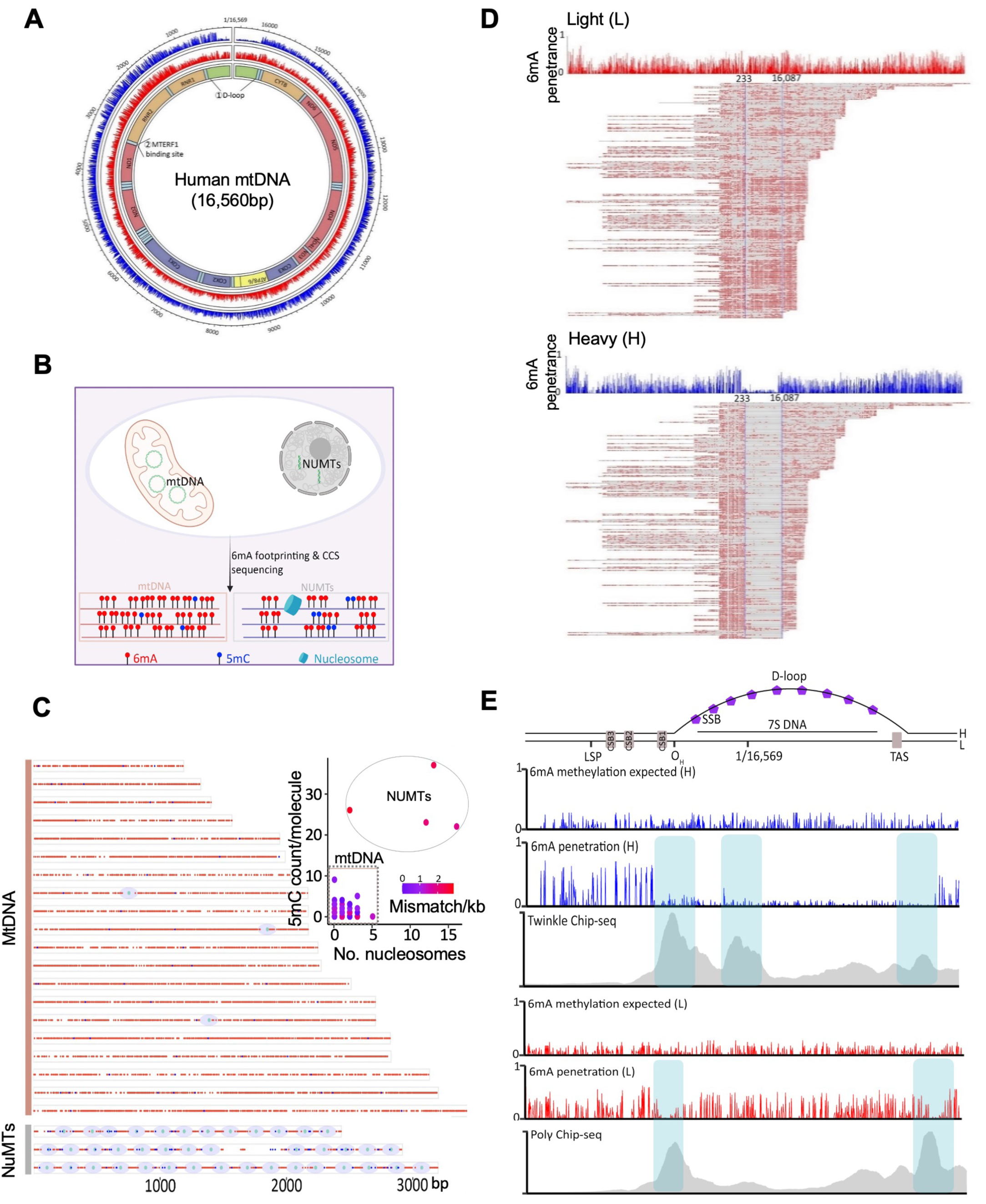
Nucleoid organization of mitochondrial DNA (mtDNA) A. Aggregated 6mA distribution on human mtDNA. The mtDNA schematic is shown as the inner circle. 6mA distribution on the heavy strand (blue) and the light strand (red) is plotted separately. B. A schematic diagram showing distinguishing features of mtDNA and NUMT detected by 6mA-FP. C. Representative mtDNA and NUMT DNA molecules detected by 6mA-FP. Purple oval, a nucleosome called by the presence of a large 6mA footprint; blue dot, 5mC; red dot, 6mA (red). Inset, the scatter plot showing the number of called nucleosomes vs. called 5mC on individual molecules (right). Color scale indicates the number of mismatched positions according to the reference mitochondrial sequence. mtDNA and NUMT molecules can be clearly distinguished. D. 6mA distributions on the light (red) and heavy strand (blue) spanning the D-loop. Bottom, individual DNA molecules sorted by the right position aligned to the reference sequence. Top, aggregated 6mA distributions. E. Fine structure of the D-loop for the light (red) and heavy strand (blue). Top, schematic for the D-loop and nearby genomic landmarks. Line plots show the observed 6mA methylation profiles as well as expected for random methylation. 6mA profile for the heavy strand (blue) and light strand (red). ChIP-seq tracks for Polγ and Twinkle are also shown, and their peak regions aligned to respective 6mA footprints are highlighted.

Human mtDNA is packaged into nucleoids with various mitochondria-specific nucleoproteins, especially TFAM [76]. In our 6mA-FP experiments with human cells, we recovered numerous mtDNA reads. We tested both Hia5 and M.EcoGII and determined that M.EcoGII was effective at methylating mtDNA, even after short incubation (Additional file 1: Fig. S13). With M.EcoGII, we found high resolution footprints for the mitochondrial transcription termination factor MTERF1 in most DNA molecules mapped to its binding site (Additional file 1: Fig. S13). Intriguingly, while 6mA levels on the heavy or light strand were similar for DNA molecules from the coding region, disparate 6mA levels on these two strands were found for the displacement loop (D-loop) within the noncoding region of mtDNA (Fig. 7D). The D-loop contains a third strand of DNA ∼650 bp in length, duplexing with the light strand (double-stranded DNA/dsDNA) and displacing the heavy strand (single-stranded DNA/ssDNA) (Fig. 7E). The strong methylation bias in favor of the light strands was detected in samples treated by M.EcoGII as well as Hia5 (Additional file 1: Fig. S13), most likely attributable to protection of the displaced heavy strand from 6mA deposition by the mitochondrial ssDNA binding protein (mtSSB) [43, 76, 77]. The unambiguous detection of the mtDNA D-loop demonstrates 6mA-FP’s power to resolve large non-canonical DNA structures and opens a new avenue for studying D-loops, R-loops, and various DNA recombination and DNA repair intermediates at the single molecule level.

The D-loop was flanked on both sides by sharp boundaries of very low 6mA densities. These 6mA footprints, present on both the heavy and light strand, aligned with ChIP-seq peaks of the mitochondrial-specific DNA helicase TWINKLE and DNA polymerase γ (POLγ) (Fig. 7E). TWINKLE is stably associated with the heavy strand [78, 79]; its footprints on the heavy strand were sharply bounded on one side by high 6mA density regions outside the D-loop and more loosely defined on the other by low 6mA density regions inside the D-loop (Fig. 7E). POLγ is stably associated with the light strand [80]; its footprints on the light strand were sharply bounded on both sides by high 6mA density regions within and without the D-loop (Fig. 7E). Importantly, two sets of POLγ footprints were unambiguously identified on both sides of the D-loop for almost all DNA molecules. These results strongly support that two sets of the mitochondrial replication machinery are loaded onto the D-loop, and they are stably arrested at the heavy strand replication origin (O_H_) and the termination-associated sequence (TAS), respectively. This conclusion can only be rigorously derived from our single-molecule long-read 6mA-FP data, but not from the previously published ensemble level short-read ChIP-seq data, highlighting 6mA-FP’s power in establishing coordinate binding events at the single molecule level.

## Discussion

### Calling base modifications with SMRT CCS

As a sequencing by synthesis technology, SMRT/PacBio sequencing takes full advantage of DNA polymerase Φ29’s high fidelity, robust strand displacement activity, and strong processivity to generate multiple strand-specific readouts of the same DNA molecule. While SMRT sequencing can generate high quality sequencing information with only a few passes (≥5 passes for each strand), it is more challenging to call base modifications, which are distinguished from their unmodified counterparts by polymerase kinetics and often present in a strand-specific manner. SMRT CCS detects 6mA by registering its kinetic signature of significantly increased IPD, relative to an unmodified A of the same sequence context (i.e., increased IPDr). However, multiple passes are needed for precise evaluation of IPD, as each subread IPD readout represents stochastic sampling of IPD values distributed in an exponential decay curve. Kinetic analysis is further complicated by random and prolonged DNA polymerase pausing. Reducing the number of passes dramatically increases the probability of overestimating IPD and miscalling 6mA. We therefore have set a high threshold (≥10 passes for each strand) for 6mA calling and optimized DNA fragment sizes (3-5kb) in SMRT CCS libraries. This threshold also allows us to add a local outlier trimming step in IPD conversion to counter the effect of polymerase pausing, by removing the top 10% of subread IPD values for each position in a DNA strand before averaging the rest to output a CCS IPD value. For the Sequel data, our new computational pipeline, ipdTrimming, matches or beats 6mA calling performance of our previous ipdSummary pipeline, while drastically reducing the computational cost. Based on our study, we postulate that adapting ipdTrimming for the Revio system may improves its 6mA calling performance, without incurring excessive computational cost.

Importantly, we have systematically compared our ipdTrimming pipeline with fibertools, the latest 6mA calling pipeline specially tailored for 6mA-FP/fiber-seq [33]. In datasets with 6mA levels ranging from <1% to approaching 100%, ipdTrimming substantially outperforms fibertools in both recall and precision. Critically, fibertools exhibits strong systematic biases: on the one hand, it frequently makes false 6mA calls at sites proximal to genuine 6mA sites, leading to over-expansion of 6mA clusters and blurring of 6mA footprints; on the other, it often misses genuine 6mA sites that are sparsely deposited due to nucleosome dynamics (both terminal DNA unwrapping and internal breathing) or other transient increase in accessibility. Fibertools results therefore severely underestimate and obscure the dynamic and heterogeneous nature of chromatin. Since Fibertools is trained on Hia5 dataset, its performance deteriorates for 6mA-FP using other MTases (e.g., M.EcoGII). Most critically, fibertools has especially poor performance in detecting endogenous 6mA at physiologically relevant levels (<1%). In contrast, our ipdTrimming pipeline excels under these conditions, offering a robust and accurate solution for 6mA detection, regardless of methylation density or chromatin dynamics. This makes ipdTrimming the optimal choice for cutting-edge epigenetic studies.

Our analysis is mainly focused on 6mA’s primary onsite effect on IPD, directly attributable to the kinetic barrier for the incoming dTTP to basepair with 6mA on the template DNA strand. Nonetheless, 6mA also has secondary offsite effects, as the template DNA strand makes extensive contact with DNA polymerase Φ29 and the *N*^6^-methyl group may indirectly affect IPD at nearby sites [36, 81]. With high 6mA density, as is the case in 6mA-FP, its secondary offsite effects are likely to increase. 6mA’s secondary offsite effects are most obvious at positions 5 nucleotides upstream (5’) of primary 6mA sites [36]. Increased IPD values at secondary sites, if an adenine, may cause their misidentification as 6mA. Intriguingly, ipdTrimming makes substantially fewer 6mA calls at one of the likely secondary sites than ipdSummary. The local trimming of IPD outliers may therefore also limit 6mA’s secondary offsite effects, contributing to ipdTrimming’s better performance with 6mA-FP data. Crosstalk between nearby base modifications (6mA and 5mC), mediated by their secondary offsite effects, complicate their calling. In this study, we use native genomic DNA instead of WGA as a negative control for 6mA-FP of human chromatin, so that the kinetics baseline will include perturbations to IPD caused by the endogenous 5mC. High-quality ground truth datasets, with synthetically defined base modifications and their combinations in a wide range of sequence context, are needed to fully resolve this difficult issue.

### Revealing epigenetic heterogeneity with 6mA-FP

In 6mA-FP, information about nucleoprotein organization is transcoded via in vitro methylation into exogenous 6mA on individual DNA molecules, which can be simultaneously revealed with endogenous 5mC signifying the epigenetic state of DNA, as well as the primary DNA sequence. Importantly, each binding event can be registered by multiple methylation sites at the single molecule level, allowing it to be called with high confidence. Here we show that by finetuning the in vitro methylation timing, we can differentiate the TF-unbound state, the TF-bound state, and the TF-transient-off state of chromatin. Short methylation, resulting in low, unsaturated 6mA levels, can preserve footprints left by dynamic binding of CTCF and other chromatin-associated proteins. Long methylation, on the other hand, resulting in near saturation 6mA levels, can reveal the dynamic nature of these binding events and distinguish them from stable binding events as well as the unbound state. Also, by optimizing the in vitro methylation condition and enriching heavily methylated DNA molecules of proper sizes, our work generates accurate and high-resolution footprints that allow for unambiguous detection of the mtDNA D-loop and the strand-specific and coordinate loading of the mitochondrial replication machinery. This represents a great improvement from previous report of the nucleoid organization of human mtDNA [76]. It also shows that 6mA-FP can be used to resolve large non-canonical DNA structures and establish joint and strand-specific binding events at the single molecule level, both of which cannot be achieved by the ensemble level short-read ChIP-seq data 6mA-FP can provide near-basepair resolution, much higher than what 5mC-based NOME-seq can achieve: on average, an adenine site is present for every 2 bp, while a GpC dinucleotide, targeted by the methyltransferase M.CviPI, is present at a much lower density (>20 bp) [13, 82, 83]. With our optimized experimental and computational framework, we have extensively mapped DNA binding events around TSS at the single-molecule level, revealing critical features of the transcription-associated epigenetic landscape at unprecedented resolution in human cells. Critically, small 6mA footprints (<50 bp) are highly enriched near active TSS, while nucleosome footprints are depleted. While binding of TFs, including CTCF, certainly contributes to these small 6mA footprints, a more intriguing scenario is stably bound Pol II, especially the preinitiation complex (PIC) and promoter proximal pausing complex (PPP), as recently reported in a fiber-seq analysis of transcription in *Drosophila* [72]. Sub-nucleosome particles (tetramer and hexamer), either standalone or in association with Pol II or chromatin remodelers, represent another alternative scenario [84]. However, we would like to point out that it is still a challenge to distinguish various binding events at the single molecule level, due to overlapping signals, background noise, and lack of specific recognition. Deeper understanding of the transcription-associated chromatin landscape at the single molecule level, especially over the range spanning transcription regulatory elements and transcriptional units, will provide critical insights into transcriptional regulation.

SMRT CCS is ideally positioned to provide the long-range, multimodal readout of genetic and epigenetic information at the single molecule level. To ascertain the overall occupancy and dynamic behavior of a nucleoprotein, many DNA molecules covering the same binding site need to be examined. This demand for high sequencing depth is a challenge to meet for mammalian systems with large nuclear genomes. Only for the high copy number rDNA and mtDNA, the heterogeneity of bound and unbound DNA molecules is fully revealed, even though it is likely a general phenomenon across the entire genome. The situation is improved by the greatly expanded capacity of the Revio system. Reduced representation, analogous to Reduced Representation Bisulfite Sequencing (RRBS) for 5mC profiling [31, 85], and target sequence enrichment, such as CRISPR-CATCH [86], promise to further increase coverage for genomic regions of interest.

## Declarations

This study was conducted in accordance with University of Southern California ethical standards. The datasets generated and/or analyzed during the current study are available with access codes in Method section. All authors consent to the publication of this manuscript in Genome Biology. A.M.W is a scientist working for Pacific Biosciences, Inc. All other authors declare no competing interests.

## Author contributions

W.T.Y. developed the computation pipeline, performed most analysis, and prepared the manuscript; X.Q.W and J.W. performed the wet-lab experiments and contributed to data interpretation; A. M. W. helped with development of the computation method; F.W helped with manuscript preparation; Y.L and Y.D. conceptualized, designed the study, and wrote the manuscript. All authors contributed to writing and revising the manuscript and approved the final version.

## Acknowledgements

We would like to acknowledge Dr. Sheng Li for her critical reading and discussion of the manuscript. This research was supported by Norris Comprehensive Cancer Center NCI P30 grant (P30 CA014089-45) and the Jane Anne Nohl Division of Hematology and Center for the Study of Blood Diseases, Keck School of Medicine to Y.D. The funding body had no role in the design of the study, collection, analysis, and interpretation of data, or in writing the manuscript.

## Methods

### SMRT CCS data processing

#### ipdSummary

Subread level SMRT CCS data were processed as previously described [32]. Briefly, each DNA molecule in the raw data was partitioned into a separate bam file using pysam module [87]. Circular Consensus Sequence (CCS) was calculated for each DNA molecule using the CCS module [34]. Only CCS reads with high passes (≥10 passes for each strand) were retained for the following analysis. Single molecule aligned BAM files were generated using BLASR [88], which in turn served as the input for the ipdSummary module to calculate IPD ratios (IPDr) [34]. DNA molecules with global or local anomalies in IPDr were removed to further improve 6mA calling accuracy.

#### ipdTrimming

Local trimming of IPD outliers at the subread level was introduced into IPD conversion: for each site in a DNA molecule, the top 10% of subread IPD values were removed; the remaining subread IPD values were then averaged to generate the CCS IPD value. This was implemented by a modified CCS module (https://gitfront.io/r/user-1129035/ActyN7zZ8hAG/6mA-footprint/tree/ccs%28ipdTrimming%29/bin/), which took in the subread level raw data and output the CCS level IPD values aligned to its own sequence. This was followed by movie-time normalization: to compensate for variations in the polymerase elongation rate on individual DNA molecules, each CCS IPD value was normalized against the CCS IPD value averaged across all sites in a DNA molecule. This CCS IPD value was then compared to that of an unmodified base with the same sequence context, provided by a kmer model pretrained on the Sequel data (https://github.com/PacificBiosciences/kineticsTools/tree/master/kineticsTools/resources). The last two steps were implemented by a custom script (ipdRatiocalculator_fromCCS.py, https://gitfront.io/r/user-1129035/ActyN7zZ8hAG/6mA-footprint/), which output a BAM file for DNA sequence tagged with kinetics parameters, including IPD ratios (IPDr).

#### Revio Data processing

The on-instrument IPD conversion for the Revio system only outputs the CCS level IPD, by averaging all subread IPD values for each site of a DNA molecule. We adapted a kmer based pipeline for 6mA calling on the Revio system. Specifically, we trained a new kmer model using the Revio data. We began by processing two input datasets: Revio-generated SMRT CCS results from 6mA-FP of human chromatin (6mA-FP HiFi.bam) and the native human genomic DNA (gDNA HiFi.bam). After movie-time normalization, gDNA HiFi.bam reads were aligned to the human reference genome (hg19) using pbmm2, generating reference IPD values for each genomic position. We then trained a new kmer model based on the Revio-sequenced human genome. Specifically, CCS IPD values across all reads covering a certain adenine in the genome (sequencing coverage depth ≥10×) were averaged to assign a reference/baseline IPD value to this genomic position. Genomic positions sharing the same local sequence context ([-10, +4]: NNNNANNNNNNNNNN; the same as the kmer model pretrained on the Sequel data) were subsequently binned, and their IPD values were averaged to finetune the new kmer model. To calculate IPDr, IPD values from 6mA-FP were compared with reference IPD values of the equivalent genomic position or kmer sequence context. We made three sets of comparisons: ***1)*** to the kmer model pretrained on the Sequel data, ***2)*** to reference IPD values assigned to the human genome, and ***3)*** to the new kmer model trained on the Sequel data.

#### 6mA calling

We first set an IPDr threshold that recalled 98.5% of GATC sites in the dam^+^ plasmid DNA positive control dataset, and then applied this threshold in the WGA negative control dataset to calculate the background noise level for 6mA calling. An alternative approach, by comparing the *Tetrahymena* native genomic DNA and WGA datasets, led to the same conclusion that the background noise level is much higher for the kmer pipeline (Additional file 1: Fig. S2). In samples with only a fraction of adenine sites methylated, their IPDr values exhibited a bimodal distribution. After logarithmic transformation, the 6mA peak with higher IPDr values was closely fitted by a Gaussian distribution curve. Gaussian demixing was performed using a custom script (m6A_gaussianmix.py), which determined an IPDr threshold above which 6mA was called, as well as the false positive and false negative rates for 6mA calling.

#### 5mC calling

5mC was called by PacBio’s primrose software with default parameters [89]. Methylation probabilities for each CpG site within the single molecule, present in the ML and MM tags of the bam file, were converted into the continuous probabilities (ranging from 0 to 1) using extract5mCforprimrose.py. CpG sites with methylation probabilities ≥0.9 were preserved for downstream analysis.

### 6mA cluster and 6mA footprint

6mA sites detected at the single-molecule level were grouped into 6mA clusters. Specifically, all 6mA sites within 15 or 30 bp of each other were combined, and multiple combined 6mA sites (≥3) formed a 6mA cluster, the size of which was the distance between the first and the last 6mA sites. Following 6mA cluster identification, 6mA footprints were determined by analyzing regions between adjacent 6mA clusters. These inter-cluster regions, if containing multiple unmodified A sites (≥3), formed a 6mA footprint, the size of which was the distance between adjacent 6mA clusters. Two alternative thresholds for combining adjacent 6mA sites into a 6mA cluster were used: the higher threshold (inter-6mA distance ≤30 bp) robustly detected large 6mA footprints corresponding to nucleosomes; the lower threshold (inter-6mA distance ≤15 bp) was more sensitive at detecting small 6mA footprints such as CTCF and Pol II, at the cost of higher background noise.

### 6mA autocorrelation analysis and classification by spectral clustering

For 6mA-FP analysis, we first filtered out DNA molecules with few 6mA sites (<30 6mA sites per kb). For autocorrelation analysis of 6mA sites, a vector consisting of a series of 0’s (no 6mA) and 1’s (6mA on either strand) was constructed for each DNA molecule. This vector was processed by the acf function within the statsmodel [90] python package to generate the full-length auto-correlogram. The head 1000 bp of the auto-correlogram for each DNA molecule was used for spectral clustering in the sklearn package [48]. The number of clusters was manually adjusted to avoid generating large numbers of clusters containing few DNA molecules.

We evaluated the regularity and period of 6mA distribution in each DNA molecule. For each DNA molecule, a vector consisting of a series of 0’s (unmodified A on either strand), NaN (C/G) and 1’s (6mA) was constructed. A 33 bp NaN-sensitive rolling mean was applied to the raw vector, enhancing signal smoothness using the R imputeTS package [91]. Autocorrelation analysis was then performed on the smoothed vector. Peaks (maximum and minimum) were identified by the find_peaks function in the python scipy module [92]. The absolute σ values (autocorrelation coefficient) of the first 8 peaks were summed up as an indicator for the regularity of 6mA distribution. The average inter-peak distance, obtained by linear regression of peak positions, was an estimate of the period of 6mA distribution, corresponding to the nucleosome repeat length (NRL) on individual chromatin fibers.

### Analysis of centromeric/peri-centromeric reads

CCS reads were aligned to the human T2T-CHM13v1.0 [93] reference genome using blastn [94] to selectively enrich the (peri)centromeric reads in silico. Using the latest annotation of the centromeric regions in the T2T genome [51], molecules fully contained within these regions were retained. For the centromeric higher-order repeat (HOR) reads, molecules were represented as a series of monomers as previously described [95]. Briefly, a catalog of distinct 58 monomers were mapped back to the long read with these following parameters: -max_target_seqs 1000000 -word_size 7 -qcov_hsp_perc 60. The assignment of monomer to a long read was calculated via dynamic programming to ensure that each segment of the read be assigned at most one monomer. All code used for these analyses is derived from the GitHub link (https://github.com/hacone/hc_temporary/releases/tag/submission-v.1.0). Importantly, a conserved GATC site in J1 allowed us to enrich DNA molecules comprising J1/J2 tandem arrays after 6mA-FP and DpnI digestion. It probably also promotes nucleosome positioning near both termini of J1/J2-containing DNA fragments, but not internally.

### Transcription factor binding analysis

Transcription factor binding motifs in the MEME format, containing a collection of position weight matrices, were downloaded from the JASPAR CORE database [96], in which a curated, non-redundant set of profiles were experimentally defined. These motif sequences were aligned to the reference genome using FIMO [97] of the MEME-Suite. Assessment of genomic binding associated with these motifs was measured by the FIMO score. Notably, a threshold of score ≥ 45, corresponding to the p-value ≤3.17e-05, was employed to identify significant matches for subsequent analysis. In parallel, CCS reads were mapped back to the same reference sequence by blastn. Only CCS reads with a mapped coverage of ≥98% and a mapped identity of ≥95% were retained for further analysis.

Bedtools [98] was utilized to find DNA molecules overlapping with these transcription factor binding sites. TF binding sites were extended by 50 bp in both upstream and downstream directions. Reads that fully covered this extended region were extracted. By combing the coordinates of reads mapped to the genome, methylation sites on single molecule and the strand information of TF binding sites, normalized methylation profile was obtained by calculating the column mean within a window of around 100 bp, centered at each TF binding site.

### Transcription start site metaplots

Transcription start sites used in this study were derived from the list of all known RefSeq transcripts in hg19 provided by HOMER v4.11.1 (http://homer.ucsd.edu/homer/data/genomes/hg19.v6.4.zip). For each protein-coding gene, the TSS of this gene is determined with these following criteria: If multiple transcripts originating from the gene share a common TSS, the TSS with the highest occurrence frequency is designated as the TSS for this gene; If no such dominant TSS exists, the midpoint of the TSSs across multiple transcripts is chosen as the gene’s TSS. The TSS for each protein-coding gene was then elongated by 50 bp in both upstream and downstream directions. TSS regions were classified based on whether these regions overlap with a specific histone modification peak by at least 1 bp. In addition, the methylation profile around TSSs was calculated in an identical manner to that described above for TF sites. CCS reads were required to fully cover the extended TSS region. Finally, CCS reads were sorted by either the 6mA cluster size overlapping with these extended regions or the expression values of these TSS-associated genes for visualization.

### Mapping CCS to mitochondrial DNA (mtDNA)

The complete circular human mitochondrial genome was downloaded from the NCBI GenBank database with the accession number NC_012920.1. To make the long reads spanning the start/end of the circular genome mappable, a linearized mitochondrial genome was generated by concatenating two tandem copies of human mtDNA. CCS reads were aligned to the modified reference using blast with the following parameters: -word_size 50, -outfmt 5. The XML format of blastn output was converted into a SAM file with the reference sequence by Blast2Bam (https://github.com/guyduche/Blast2Bam). Reads with ≤98% mapping coverage or ≤95% mapping identity were excluded. Since the two identical mitochondrial genome sequences were joined, in principle, reads than don’t span the start/end region could be potentially aligned to two separate regions. To ensure unique mapping for each read, only reads aligned to the first one (with small reference coordinates) of these two regions were retained. In the SAM file, CIGAR values describe how the read aligns to the reference genome, including insertions, deletions, mismatches and matches, allowing for the alignment at the single nucleotide resolution. By parsing the CIGAR values in the SAM file, methylation sites on both strands of individual DNA molecules were mapped back to the mitochondrial genome.

### Mapping CCS to ribosomal DNA (rDNA)

In the context of the hg19 genome assembly, a single repeat unit sequence (NCBI U13369.1) rDNA was inserted into chr13. To enhance the mappability of the long reads derived from the rDNA, a modified reference rDNA was created. This modified sequence consists of intergenic sequence (IGS), 45s coding rDNA and IGS, effectively representing one and a half tandem copies of rDNA. CCS reads mapping was processed in an identical manner to that described for mtDNA. Briefly, reads were mapped using blastn and then converted into the SAM file. Low alignment quality reads (≤98% mapping coverage or ≤95% mapping identity) were removed. Reads mappable to both IGS regions were allocated to the first.

### Cell culture

Human acute myeloid leukemia (AML) cell lines OCI-AML3 (DSMZ no. ACC 582, obtained from Dr. Xiaotian Zhang at University of Texas Health Science Center at Houston) and MOLM13 (DSMZ no. 554) were cultured in RPMI-1640 (GIBCO™ 11875-119) supplemented with 10% fetal bovine serum (Sigma) and 1% penicillin/streptomycin (GIBCO™ 15140-163).

### Sample DNA preparation

A *Pvu*I-digested fragment of the pBluescript II SK(-) plasmid (∼1kb) purified from dam^+^ *E coli* was used as the positive control sample for 6mA calling. Native genomic DNA was extracted from *Tetrahymena* WT cells (SB210, *Tetrahymena* Stock Center) using Wizard® Genomic DNA Purification Kit (Promega). REPLI-g Single Cell Kit (Qiagen) was used for whole genome amplification (WGA) to generate the negative control sample. DNA samples were sheared to 3-5kb in length with Megaruptor (Diagenode Diagnostics) and prepared for SMRT CCS.

1.7×10^5^ OCI-AML3 or MOLM13 cells were lysed in 0.5 mL of nuclear extraction buffer (20mM HEPES pH 7.9, 10mM KCl, 0.1% Triton X-100, 20% glycerol, 0.5mM spermidine, 1× Protease Inhibitor Cocktail) for 8 min on ice. Nuclei were isolated by centrifuge at 600 g for 5 minutes with two washes of nuclear extraction buffer. Purified nuclei were methylated in a 30μL of 50mM Tris-HCl pH 8.0, 2mM EDTA, 0.5mM EDGA, 160μM SAM (New England Biolabs), 1× Protease Inhibitor Cocktail and 38μM recombinant M.EcoGII or Hia5 for 1 hour (short methylation) or 3 hours (long methylation) at 37°C. Genomic DNA was extracted with Monarch® HMW DNA Extraction Kit for Cells & Blood (New England Biolabs), digested with *Dpn*I overnight at 37°C, and resolved on 1% agarose gel. DNA fragments between 3 to 5 kb were gel extracted using the QIAEX II Gel Extraction Kit (Qiagen). Extracted DNA was quantified with the Quant-iT™ dsDNA Assay Kits, high sensitivity (HS) and broad range (BR) (Invitrogen™ Q33120) on a Qubit™ 3 Fluorometer. Samples were submitted for SMRT CCS at the Genomics Core Facility at the Icahn School of Medicine at Mount Sinai.

### Protein Purification

Expression and purification of pA-M.EcoGII and pA-Hia5 were performed as previously described [99]. Briefly, bacterial culture was grown to 0.5 OD600 in LB medium containing 50 μg/mL kanamycin (Sigma) and induced overnight with 0.4 mM IPTG at 16°C. Cells were lysed in high salt buffer (1 M NaCl, 10 mM Tris-HCl (pH 7.5), 10 mM Imidazole, 0.5% NP-40, 10% glycerol, 1 mM PMSF and protease inhibitor cocktail) and sonicated. Proteins were purified on Ni-NTA beads and washed in low-salt buffer (250 mM NaCl, 10 mM Tris-HCl (pH7.5), 20 mM Imidazole, 0.5% NP-40, 10% glycerol, 1 mM PMSF and protease inhibitor cocktail) and eluted (250 mM Imidazole). Eluate was dialyzed overnight with ULP1, and the cleaved tag was removed by Ni-NTA binding.

### Availability of data and materials

Raw sequencing data generated in this study have been deposited in the NCBI under accession numbers PRJNA1166413, which can been viewed with this link: https://dataview.ncbi.nlm.nih.gov/object/PRJNA1166413?reviewer=dboh1jkhoprtplbbt0rg8tsca5.

All code used in this study is available at https://gitfront.io/r/user-1129035/ActyN7zZ8hAG/6mA-footprint/. Previously reported PacBio sequencing data, including *Tetrahymena* native genome DNA (SRR16944761), *Tetrahymena* whole genome amplification (SRR28602303), fiber-seq in human OCI-AML3 (SRR17010848), fiber-seq in Widom 601 (SRR13173521), and a *PvuI*-digested fragment of the pBluescript II SK(-) plasmid (SRR27430828) are publicly available.

## Supplemental figure legends

**Figure S1. Comparing 6mA called by CCS-kmer, ipdSummary, and ipdTrimming in dam^+^ plasmid DNA**

A. Left schematic, palindromic GATC sites in four distinct methylation states: full methylation on both strands, hemi-methylation on either strand, or no methylation. Right schematic, three GATC sites in this fragment of plasmid DNA. Three prominent combination of methylation states: [3, 3], all three sites methylated on both the Watson and Crick strands; [3, 0], all three sites only methylated on the Watson strand; [0, 3], all three sites only methylated on the Crick strand.
B. ipdTrimming improves 6mA calling for a representative position as compared to ipdSummary.
C. IPDr distribution in CCS-kmer, ipdSummary, and ipdTrimming. A log_2_(IPDr) threshold of 0.7 was set for calling 6mA in all three pipelines.
D. Table comparing all combinations of methylation states for the three GATC sites called by CCS-kmer, ipdSummary, and ipdTrimming.

**Figure S2. Comparing 6mA called by CCS-kmer, ipdSummary, and ipdTrimming in native *Tetrahymena* genomic DNA**

A. IPDr distribution for all A sites (blue) vs. A sites in the ApT dinucleotide (red), by CCS-kmer, ipdSummary, and ipdTrimming.
B. Differential between the A curve and the ApT curve (see above). Residue levels, a reflection of 6mA calling background noise in *Tetrahymena* genomic DNA, are shown for CCS-kmer, ipdSummary, and ipdTrimming, respectively.
C. 6mA calling background noise levels, estimated by comparing native DNA (blue) and WGA DNA (red) with CCS-kmer, ipdSummary, and ipdTrimming.
D. Venn diagram for 6mA sites called by ipdSummary and ipdTrimming. Overlap 6mA sites, representing the overwhelming majority, are shown at the center. Unique 6mA sites are shown at the top or bottom.
E. 2D-distribution of the four methylation states of ApT dinucleotides detected by ipdSummary or ipdTrimming as indicated on top. X and Y axes correspond to the log_2_(IPDr) of A sites on Watson and Crick strands, respectively.
F. Compare ipdSummary and ipdTrimming for detection of segregation strand bias of hemi-6mApT in hemi^+^ DNA molecules. X-axis: segregation strand bias for hemi-6mApT, defined as the difference-sum ratio between hemi-W and hemi-C: 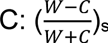. Y-axis: the percentage of hemi^+^ molecules with segregation strand bias specified on X-axis.

**Figure S3. Comparing 6mA called by ipdSummary and ipdTrimming in 6mA-FP of the Widom 601 nucleosome array**

A. Gaussian demixing of IPDr distribution for ipdSummary and ipdTrimming.
B. Table showing key statistics for 6mA called by ipdSummary and ipdTrimming.
C. 6mA distribution across the entire length of the Widom 601 nucleosome array (X-axis). Y-axis: the count of joint and differential 6mA calls.
D. Enlarged view of panel C, focusing on one nucleosome-linker DNA region to show joint and differential 6mA calls.

**Figure S4. Comparing 6mA called by fibertools and ipdTrimming**

A. Representative *Tetrahymena* genomic region containing 6mA sites called by ipdTrimming and fibertools, respectively. Note that ipdTrimming only called 6mA on the ApT dinucleotide, while fibertools called many 6mA on non-ApT dinucleotides.
B. 6mA calling on the ApT dinucleotide. Fibertools showed improved recall rates and FPR under this condition.
C. For all positions on the Widom 601 nucleosome array, compare the number of 6mA sites called by fibertools, ipdTrimming, and ipdSummary. Note that ipdTrimming and ipdSummary closely matched each other, while fibertools diverged substantially.
D. Top, 6mA distribution across the entire length of the Widom 601 nucleosome array (X-axis). Y-axis: the count of joint and differential 6mA calls. Bottom, enlarged view focusing on one nucleosome-linker DNA region to show joint and differential 6mA calls.

**Figure S5. Comparing 6mA called by all four methods in 6mA-FP of human chromatin**

**Figure S6. The computation pipeline for 6mA calling using the Revio data.**

**Figure S7. Optimization of 6mA-FP procedure**

A. Schematic for mapping distribution of chromatin-associated proteins by 6mA-FP.
B. Methylation probability for all 16 NAN combinations. In vitro methylation of human chromatin by M.EcoGII showed less intrinsic sequence bias than Hia5.
C. Aggregated 6mA distribution around CTCF-high binding sites from 6mA-FP of human chromatin. Note the higher signal-to-noise ratio in M.EcoGII than in Hia5.
D. Size selected genomic DNA (5kb) from OCI-AML3 or MOLM13 cells after short (1hr) or long (3hr) methylation treatment by M.EcoGII. DNA after DpnI digestion was shown on the right.

**Figure S8. Centromeric chromatin organization revealed by 6mA-FP**

A. Schematic for human centromere organization.
B. Diversity in 6mA patterns of centromeric heterochromatin. Percentage of DNA molecules in the five classes (as defined in Fig. 3A) are shown for each centromeric region.
C. 6mA distribution period, corresponding to NRL, in DNA molecules mapped to each centromeric region.
D. Distribution of all centromeric DNA molecules across the five classes.

**Figure S9. Detection of CTCF binding at the single molecule level**

A. Top, aggregated distribution of 5mC around Rank 1 (CTCF-high, red) and Rank 5 (CTCF-low, blue) CTCF binding sites. Bottom, 6mA sites on individual molecule. Human chromatin was treated for short methylation (M.EcoGII, 1 hour).
B. The same setup, but for human chromatin treated for long methylation (M.EcoGII, 3 hour).
C. 6mA clusters around CTCF binding motifs revealed by short methylation 6mA-FP. DNA molecules are ranked as in Fig. 4B. 6mA clusters are color coded by the size, as in Fig. 4F.
D. Heatmap for distribution of size (y-axis) and position (x-axis, relative to the CTCF binding motif) of 6mA clusters in CTCF-high (Rank 1) and CTCF-low (Rank 5) DNA molecules. DNA molecules are ranked as in Fig. 4B.

**Figure S10. Representative genomic locus in MOLM13 cells.**

A. A representative locus (at the gene promoter of WRAP73) enriched for large 6mA clusters (top tracks) and TFs in MOLM13 cells.
B. Large 6mA clusters are preferentially flanked by active histone marks, but not repressive ones.

**Figure S11. SPI1/PU.1 and CEBPα binding in rDNA**

A. 6mA (left) and 5mC sites (right) on DNA molecules mapped to the SPI1/PU.1 binding site in rDNA from OCI-AML3 and MOLM13 cells, respectively.
B. 6mA (left) and 5mC sites (right) on DNA molecules mapped to the CEBPα binding site in rDNA from OCI-AML3 and MOLM13 cells, respectively.

**Figure S12. The 5mC level shows the opposite trend to the 6mA level at TSS.**

Aggregated distribution of 5mC around TSS. The TSS are grouped based on histone modifications status (**A**) or gene expression level (**B**) as indicated on top.

**Figure S13. 6mA-FP of mitochondrial nucleoid.**

A. Comparing 6mA-FP results by M.EcoGII vs. Hia5.
B. MTERF1 footprint, revealed by M.EcoGII 6mA-FP.
C. 6mA distribution around the D-loop, revealed by Hia5 6mA-FP.

## Supplemental tables

**Table S1. SMRT CCS datasets used in this work.**

**Table S2. Statistics for Gaussian demixing of 6mA and unmodified A peaks.**

**Table S3. 6mA calling statistics for fiber-seq data.**

## References

1. Rivera CM, Ren B: Mapping human epigenomes. Cell 2013, 155:39–55.

2. Mehrmohamadi M, Sepehri MH, Nazer N, Norouzi MR: A comparative overview of epigenomic profiling methods. Front Cell Dev Biol 2021, 9:714687.

3. Johnson DS, Mortazavi A, Myers RM, Wold B: Genome-wide mapping of in vivo protein-DNA interactions. Science 2007, 316:1497–1502.

4. Mikkelsen TS, Ku M, Jaffe DB, Issac B, Lieberman E, Giannoukos G, Alvarez P, Brockman W, Kim TK, Koche RP, et al: Genome-wide maps of chromatin state in pluripotent and lineage-committed cells. Nature 2007, 448:553–560.

5. Robertson G, Hirst M, Bainbridge M, Bilenky M, Zhao Y, Zeng T, Euskirchen G, Bernier B, Varhol R, Delaney A, et al: Genome-wide profiles of STAT1 DNA association using chromatin immunoprecipitation and massively parallel sequencing. Nat Methods 2007, 4:651–657.

6. Barski A, Cuddapah S, Cui K, Roh TY, Schones DE, Wang Z, Wei G, Chepelev I, Zhao K: High-resolution profiling of histone methylations in the human genome. Cell 2007, 129:823–837.

7. Buenrostro JD, Giresi PG, Zaba LC, Chang HY, Greenleaf WJ: Transposition of native chromatin for fast and sensitive epigenomic profiling of open chromatin, DNA-binding proteins and nucleosome position. Nat Methods 2013, 10:1213–1218.

8. Zaret K: Micrococcal nuclease analysis of chromatin structure. Curr Protoc Mol Biol 2005, Chapter 21:Unit 21. 21.

9. Song L, Crawford GE: DNase-seq: a high-resolution technique for mapping active gene regulatory elements across the genome from mammalian cells. Cold Spring Harb Protoc 2010, 2010:pdb prot5384.

10. Skene PJ, Henikoff S: An efficient targeted nuclease strategy for high-resolution mapping of DNA binding sites. Elife 2017, 6.

11. Kaya-Okur HS, Wu SJ, Codomo CA, Pledger ES, Bryson TD, Henikoff JG, Ahmad K, Henikoff S: CUT&Tag for efficient epigenomic profiling of small samples and single cells. Nat Commun 2019, 10:1930.

12. van Steensel B, Henikoff S: Identification of in vivo DNA targets of chromatin proteins using tethered dam methyltransferase. Nat Biotechnol 2000, 18:424–428.

13. Kelly TK, Liu Y, Lay FD, Liang G, Berman BP, Jones PA: Genome-wide mapping of nucleosome positioning and DNA methylation within individual DNA molecules. Genome Res 2012, 22:2497–2506.

14. Stergachis AB, Debo BM, Haugen E, Churchman LS, Stamatoyannopoulos JA: Single-molecule regulatory architectures captured by chromatin fiber sequencing. Science 2020, 368:1449–1454.

15. Shipony Z, Marinov GK, Swaffer MP, Sinnott-Armstrong NA, Skotheim JM, Kundaje A, Greenleaf WJ: Long-range single-molecule mapping of chromatin accessibility in eukaryotes. Nat Methods 2020, 17:319–327.

16. Abdulhay NJ, McNally CP, Hsieh LJ, Kasinathan S, Keith A, Estes LS, Karimzadeh M, Underwood JG, Goodarzi H, Narlikar GJ, Ramani V: Massively multiplex single-molecule oligonucleosome footprinting. eLife 2020, 9:e59404.

17. Altemose N, Maslan A, Smith OK, Sundararajan K, Brown RR, Detweiler AM, Neff N, Miga KH, Straight AF, Streets A: DiMeLo-seq: a long-read, single-molecule method for mapping protein-DNA interactions genome-wide. Nat Methods 2022, 19:711–723.

18. Wenger AM, Peluso P, Rowell WJ, Chang PC, Hall RJ, Concepcion GT, Ebler J, Fungtammasan A, Kolesnikov A, Olson ND, et al: Accurate circular consensus long-read sequencing improves variant detection and assembly of a human genome. Nat Biotechnol 2019, 37:1155–1162.

19. Flusberg BA, Webster DR, Lee JH, Travers KJ, Olivares EC, Clark TA, Korlach J, Turner SW: Direct detection of DNA methylation during single-molecule, real-time sequencing. Nat Methods 2010, 7:461–465.

20. Douvlataniotis K, Bensberg M, Lentini A, Gylemo B, Nestor CE: No evidence for DNA N (6)-methyladenine in mammals. Sci Adv 2020, 6:eaay3335.

21. Bochtler M, Fernandes H: DNA adenine methylation in eukaryotes: Enzymatic mark or a form of DNA damage? Bioessays 2021, 43:e2000243.

22. O’Brown ZK, Boulias K, Wang J, Wang SY, O’Brown NM, Hao Z, Shibuya H, Fady PE, Shi Y, He C, et al: Sources of artifact in measurements of 6mA and 4mC abundance in eukaryotic genomic DNA. BMC Genomics 2019, 20:445.

23. Kong Y, Cao L, Deikus G, Fan Y, Mead EA, Lai W, Zhang Y, Yong R, Sebra R, Wang H, et al: Critical assessment of DNA adenine methylation in eukaryotes using quantitative deconvolution. Science 2022, 375:515–522.

24. Nanda AS, Wu K, Irkliyenko I, Woo B, Ostrowski MS, Clugston AS, Sayles LC, Xu L, Satpathy AT, Nguyen HG, et al: Direct transposition of native DNA for sensitive multimodal single-molecule sequencing. Nat Genet 2024, 56:1300–1309.

25. Kong Y, Mead EA, Fang G: Navigating the pitfalls of mapping DNA and RNA modifications. Nat Rev Genet 2023, 24:363–381.

26. Eid J, Fehr A, Gray J, Luong K, Lyle J, Otto G, Peluso P, Rank D, Baybayan P, Bettman B, et al: Real-time DNA sequencing from single polymerase molecules. Science 2009, 323:133–138.

27. Wang Y, Chen X, Sheng Y, Liu Y, Gao S: N^6^-adenine DNA methylation is associated with the linker DNA of H2A.Z-containing well-positioned nucleosomes in Pol II-transcribed genes in *Tetrahymena*. Nucleic Acids Res 2017, 45:11594–11606.

28. Wang Y, Sheng Y, Liu Y, Zhang W, Cheng T, Duan L, Pan B, Qiao Y, Liu Y, Gao S: A distinct class of eukaryotic MT-A70 methyltransferases maintain symmetric DNA N^6^-adenine methylation at the ApT dinucleotides as an epigenetic mark associated with transcription. Nucleic Acids Res 2019, 47:11771–11789.

29. Beh LY, Debelouchina GT, Clay DM, Thompson RE, Lindblad KA, Hutton ER, Bracht JR, Sebra RP, Muir TW, Landweber LF: Identification of a DNA N^6^-adenine methyltransferase complex and its impact on chromatin organization. Cell 2019, 177:1781–1796.

30. Beaulaurier J, Schadt EE, Fang G: Deciphering bacterial epigenomes using modern sequencing technologies. Nat Rev Genet 2019, 20:157–172.

31. Fang G, Munera D, Friedman DI, Mandlik A, Chao MC, Banerjee O, Feng Z, Losic B, Mahajan MC, Jabado OJ, et al: Genome-wide mapping of methylated adenine residues in pathogenic Escherichia coli using single-molecule real-time sequencing. Nat Biotechnol 2012, 30:1232–1239.

32. Sheng Y, Wang Y, Yang W, Wang XQ, Lu J, Pan B, Nan B, Liu Y, Ye F, Li C, et al: Semiconservative transmission of DNA N (6)-adenine methylation in a unicellular eukaryote. Genome Res 2024, 34:740–756.

33. Jha A, Bohaczuk SC, Mao Y, Ranchalis J, Mallory BJ, Min AT, Hamm MO, Swanson E, Dubocanin D, Finkbeiner C, et al: DNA-m6A calling and integrated long-read epigenetic and genetic analysis with fibertools. Genome Res 2024:gr.279095.279124.

34. Biosciences P: SMRT® tools reference guide (v13.0). Pacific Biosciences, Menlo Park, CA 2024, https://www.pacb.com/wp-content/uploads/SMRT-Tools-Reference-Guide-v13.0.pdf:p 54.

35. Marks P, Banerjee O, Alexander D: Detection and identification of base modifications with single molecule real-time sequencing data. GitHub 2012, https://github.com/PacificBiosciences/kineticsTools/blob/master/doc/whitepaper/kinetics.pd f.

36. Clark TA, Murray IA, Morgan RD, Kislyuk AO, Spittle KE, Boitano M, Fomenkov A, Roberts RJ, Korlach J: Characterization of DNA methyltransferase specificities using single-molecule, real-time DNA sequencing. Nucleic Acids Res 2012, 40:e29.

37. Lowary PT, Widom J: New DNA sequence rules for high affinity binding to histone octamer and sequence-directed nucleosome positioning. J Mol Biol 1998, 276:19–42.

38. Davey CA, Sargent DF, Luger K, Maeder AW, Richmond TJ: Solvent mediated interactions in the structure of the nucleosome core particle at 1.9 a resolution. J Mol Biol 2002, 319:1097–1113.

39. Armeev GA, Kniazeva AS, Komarova GA, Kirpichnikov MP, Shaytan AK: Histone dynamics mediate DNA unwrapping and sliding in nucleosomes. Nat Commun 2021, 12:2387.

40. Gansen A, Felekyan S, Kuhnemuth R, Lehmann K, Toth K, Seidel CAM, Langowski J: High precision FRET studies reveal reversible transitions in nucleosomes between microseconds and minutes. Nat Commun 2018, 9:4628.

41. Chua EY, Vasudevan D, Davey GE, Wu B, Davey CA: The mechanics behind DNA sequence-dependent properties of the nucleosome. Nucleic Acids Res 2012, 40:6338–6352.

42. Vasudevan D, Chua EYD, Davey CA: Crystal structures of nucleosome core particles containing the ’601’ strong positioning sequence. J Mol Biol 2010, 403:1–10.

43. Murray IA, Morgan RD, Luyten Y, Fomenkov A, Correa IR, Jr., Dai N, Allaw MB, Zhang X, Cheng X, Roberts RJ: The non-specific adenine DNA methyltransferase M.EcoGII. Nucleic Acids Res 2018, 46:840–848.

44. Drozdz M, Piekarowicz A, Bujnicki JM, Radlinska M: Novel non-specific DNA adenine methyltransferases. Nucleic Acids Res 2012, 40:2119–2130.

45. Matsuo Y, MacLeod RA, Uphoff CC, Drexler HG, Nishizaki C, Katayama Y, Kimura G, Fujii N, Omoto E, Harada M, Orita K: Two acute monocytic leukemia (AML-M5a) cell lines (MOLM-13 and MOLM-14) with interclonal phenotypic heterogeneity showing MLL-AF9 fusion resulting from an occult chromosome insertion, ins(11;9)(q23;p22p23). Leukemia 1997, 11:1469–1477.

46. Quentmeier H, Martelli MP, Dirks WG, Bolli N, Liso A, Macleod RA, Nicoletti I, Mannucci R, Pucciarini A, Bigerna B, et al: Cell line OCI/AML3 bears exon-12 NPM gene mutation-A and cytoplasmic expression of nucleophosmin. Leukemia 2005, 19:1760–1767.

47. Baldi S, Korber P, Becker PB: Beads on a string-nucleosome array arrangements and folding of the chromatin fiber. Nat Struct Mol Biol 2020, 27:109–118.

48. Pedregosa F, Varoquaux G, Gramfort A, Michel V, Thirion B, Grisel O, Blondel M, Prettenhofer P, Weiss R, Dubourg VJtJomLr: Scikit-learn: machine learning in Python. J Mach Learn Res 2011, 12:2825–2830.

49. Lai B, Gao W, Cui K, Xie W, Tang Q, Jin W, Hu G, Ni B, Zhao K: Principles of nucleosome organization revealed by single-cell micrococcal nuclease sequencing. Nature 2018, 562:281–285.

50. Hasson D, Panchenko T, Salimian KJ, Salman MU, Sekulic N, Alonso A, Warburton PE, Black BE: The octamer is the major form of CENP-A nucleosomes at human centromeres. Nat Struct Mol Biol 2013, 20:687–695.

51. Altemose N, Logsdon GA, Bzikadze AV, Sidhwani P, Langley SA, Caldas GV, Hoyt SJ, Uralsky L, Ryabov FD, Shew CJ, et al: Complete genomic and epigenetic maps of human centromeres. Science 2022, 376:eabl4178.

52. Funk CC, Casella AM, Jung S, Richards MA, Rodriguez A, Shannon P, Donovan-Maiye R, Heavner B, Chard K, Xiao Y, et al: Atlas of transcription factor binding sites from ENCODE DNase hypersensitivity data across 27 tissue types. Cell Rep 2020, 32:108029.

53. Lambert SA, Jolma A, Campitelli LF, Das PK, Yin Y, Albu M, Chen X, Taipale J, Hughes TR, Weirauch MT: The human transcription factors. Cell 2018, 172:650–665.

54. Zhu F, Farnung L, Kaasinen E, Sahu B, Yin Y, Wei B, Dodonova SO, Nitta KR, Morgunova E, Taipale M, et al: The interaction landscape between transcription factors and the nucleosome. Nature 2018, 562:76–81.

55. Hashimoto H, Wang D, Horton JR, Zhang X, Corces VG, Cheng X: Structural basis for the versatile and methylation-dependent binding of CTCF to DNA. Mol Cell 2017, 66:711–720 e713.

56. Yang J, Horton JR, Liu B, Corces VG, Blumenthal RM, Zhang X, Cheng X: Structures of CTCF-DNA complexes including all 11 zinc fingers. Nucleic Acids Res 2023, 51:8447–8462.

57. Nakahashi H, Kieffer Kwon KR, Resch W, Vian L, Dose M, Stavreva D, Hakim O, Pruett N, Nelson S, Yamane A, et al: A genome-wide map of CTCF multivalency redefines the CTCF code. Cell Rep 2013, 3:1678–1689.

58. Huang H, Zhu Q, Jussila A, Han Y, Bintu B, Kern C, Conte M, Zhang Y, Bianco S, Chiariello AM, et al: CTCF mediates dosage-and sequence-context-dependent transcriptional insulation by forming local chromatin domains. Nat Genet 2021, 53:1064–1074.

59. Soochit W, Sleutels F, Stik G, Bartkuhn M, Basu S, Hernandez SC, Merzouk S, Vidal E, Boers R, Boers J, et al: CTCF chromatin residence time controls three-dimensional genome organization, gene expression and DNA methylation in pluripotent cells. Nat Cell Biol 2021, 23:881–893.

60. Li Y, Haarhuis JHI, Sedeno Cacciatore A, Oldenkamp R, van Ruiten MS, Willems L, Teunissen H, Muir KW, de Wit E, Rowland BD, Panne D: The structural basis for cohesin-CTCF-anchored loops. Nature 2020, 578:472–476.

61. Agarwal H, Reisser M, Wortmann C, Gebhardt JCM: Direct Observation of Cell-Cycle-Dependent Interactions between CTCF and Chromatin. Biophys J 2017, 112:2051–2055.

62. Hansen AS, Pustova I, Cattoglio C, Tjian R, Darzacq X: CTCF and cohesin regulate chromatin loop stability with distinct dynamics. eLife 2017, 6:e25776.

63. Oki S, Ohta T, Shioi G, Hatanaka H, Ogasawara O, Okuda Y, Kawaji H, Nakaki R, Sese J, Meno C: ChIP-Atlas: a data-mining suite powered by full integration of public ChIP-seq data. EMBO Rep 2018, 19:e46255.

64. Smith JS, Lappin KM, Craig SG, Liberante FG, Crean CM, McDade SS, Thompson A, Mills KI, Savage KI: Chronic loss of STAG2 leads to altered chromatin structure contributing to de-regulated transcription in AML. J Transl Med 2020, 18:339.

65. Michel BC, D’Avino AR, Cassel SH, Mashtalir N, McKenzie ZM, McBride MJ, Valencia AM, Zhou Q, Bocker M, Soares LMM, et al: A non-canonical SWI/SNF complex is a synthetic lethal target in cancers driven by BAF complex perturbation. Nat Cell Biol 2018, 20:1410–1420.

66. Antony C, George SS, Blum J, Somers P, Thorsheim CL, Wu-Corts DJ, Ai Y, Gao L, Lv K, Tremblay MG, et al: Control of ribosomal RNA synthesis by hematopoietic transcription factors. Mol Cell 2022, 82:3826–3839 e3829.

67. Heberle E, Bardet AF: Sensitivity of transcription factors to DNA methylation. Essays Biochem 2019, 63:727–741.

68. Frederick MA, Williamson KE, Fernandez Garcia M, Ferretti MB, McCarthy RL, Donahue G, Luzete Monteiro E, Takenaka N, Reynaga J, Kadoch C, Zaret KS: A pioneer factor locally opens compacted chromatin to enable targeted ATP-dependent nucleosome remodeling. Nat Struct Mol Biol 2023, 30:31–37.

69. de la Rica L, Rodriguez-Ubreva J, Garcia M, Islam AB, Urquiza JM, Hernando H, Christensen J, Helin K, Gomez-Vaquero C, Ballestar E: PU.1 target genes undergo Tet2-coupled demethylation and DNMT3b-mediated methylation in monocyte-to-osteoclast differentiation. Genome Biol 2013, 14:R99.

70. Joshi K, Liu S, Breslin SJP, Zhang J: Mechanisms that regulate the activities of TET proteins. Cell Mol Life Sci 2022, 79:363.

71. Kim S, Wysocka J: Deciphering the multi-scale, quantitative cis-regulatory code. Mol Cell 2023, 83:373–392.

72. Tullius TW, Isaac RS, Dubocanin D, Ranchalis J, Churchman LS, Stergachis AB: RNA polymerases reshape chromatin architecture and couple transcription on individual fibers. Mol Cell 2024, 84:3209–3222 e3205.

73. Falkenberg M, Larsson NG, Gustafsson CM: Replication and Transcription of Human Mitochondrial DNA. Annu Rev Biochem 2024, 93:47–77.

74. Kopinski PK, Singh LN, Zhang S, Lott MT, Wallace DC: Mitochondrial DNA variation and cancer. Nat Rev Cancer 2021, 21:431–445.

75. Wei W, Schon KR, Elgar G, Orioli A, Tanguy M, Giess A, Tischkowitz M, Caulfield MJ, Chinnery PF: Nuclear-embedded mitochondrial DNA sequences in 66,083 human genomes. Nature 2022, 611:105–114.

76. Isaac RS, Tullius TW, Hansen KG, Dubocanin D, Couvillion M, Stergachis AB, Churchman LS: Single-nucleoid architecture reveals heterogeneous packaging of mitochondrial DNA. Nat Struct Mol Biol 2024, 31:568–577.

77. Brown TA, Cecconi C, Tkachuk AN, Bustamante C, Clayton DA: Replication of mitochondrial DNA occurs by strand displacement with alternative light-strand origins, not via a strand-coupled mechanism. Genes Dev 2005, 19:2466–2476.

78. Peter B, Falkenberg M: TWINKLE and other human mitochondrial DNA helicases: structure, function and disease. Genes (Basel) 2020, 11:408.

79. Milenkovic D, Matic S, Kuhl I, Ruzzenente B, Freyer C, Jemt E, Park CB, Falkenberg M, Larsson NG: TWINKLE is an essential mitochondrial helicase required for synthesis of nascent D-loop strands and complete mtDNA replication. Hum Mol Genet 2013, 22:1983–1993.

80. Lee YS, Kennedy WD, Yin YW: Structural insight into processive human mitochondrial DNA synthesis and disease-related polymerase mutations. Cell 2009, 139:312–324.

81. Berman AJ, Kamtekar S, Goodman JL, Lazaro JM, de Vega M, Blanco L, Salas M, Steitz TA: Structures of phi29 DNA polymerase complexed with substrate: the mechanism of translocation in B-family polymerases. EMBO J 2007, 26:3494–3505.

82. Lee I, Razaghi R, Gilpatrick T, Molnar M, Gershman A, Sadowski N, Sedlazeck FJ, Hansen KD, Simpson JT, Timp W: Simultaneous profiling of chromatin accessibility and methylation on human cell lines with nanopore sequencing. Nat Methods 2020, 17:1191–1199.

83. Wang Y, Wang A, Liu Z, Thurman AL, Powers LS, Zou M, Zhao Y, Hefel A, Li Y, Zabner J, Au KF: Single-molecule long-read sequencing reveals the chromatin basis of gene expression. Genome Res 2019, 29:1329–1342.

84. Lai WKM, Pugh BF: Understanding nucleosome dynamics and their links to gene expression and DNA replication. Nat Rev Mol Cell Biol 2017, 18:548–562.

85. Meissner A, Gnirke A, Bell GW, Ramsahoye B, Lander ES, Jaenisch R: Reduced representation bisulfite sequencing for comparative high-resolution DNA methylation analysis. Nucleic Acids Res 2005, 33:5868–5877.

86. Hung KL, Luebeck J, Dehkordi SR, Colon CI, Li R, Wong IT, Coruh C, Dharanipragada P, Lomeli SH, Weiser NE, et al: Targeted profiling of human extrachromosomal DNA by CRISPR-CATCH. Nat Genet 2022, 54:1746–1754.

87. Gilman P, Janzou S, Guittet D, Freeman J, DiOrio N, Blair N, Boyd M, Neises T, Wagner M: PySAM (Python wrapper for system advisor model “SAM”) [SWR-19-57]. Computer software 2019, https://github.com/NREL/pysam.

88. Chaisson MJ, Tesler G: Mapping single molecule sequencing reads using basic local alignment with successive refinement (BLASR): application and theory. BMC bioinformatics 2012, 13:1–18.

89. Portik D, Hall R, Nyquist K, Wenger A: Extracting CpG methylation from PacBio HiFi whole genome sequencing. Pacific Biosciences, Menlo Park, CA 2022, https://www.pacb.com/wp-content/uploads/AGBT-2022-extracting-CpG-methylation-Portik-poster.pdf.

90. Seabold S, Perktold J: Statsmodels: econometric and statistical modeling with Python. SciPy 2010, 57:10–25080.

91. Moritz S, Bartz-Beielstein T: imputeTS: time series missing value imputation in R. R J 2017, 9:207.

92. Virtanen P, Gommers R, Oliphant TE, Haberland M, Reddy T, Cournapeau D, Burovski E, Peterson P, Weckesser W, Bright J: SciPy 1.0: fundamental algorithms for scientific computing in Python. Nature methods 2020, 17:261–272.

93. Nurk S, Koren S, Rhie A, Rautiainen M, Bzikadze AV, Mikheenko A, Vollger MR, Altemose N, Uralsky L, Gershman A: The complete sequence of a human genome. Science 2022, 376:44–53.

94. Ye J, McGinnis S, Madden TL: BLAST: improvements for better sequence analysis. Nucleic acids research 2006, 34:W6–W9.

95. Suzuki Y, Myers EW, Morishita S: Rapid and ongoing evolution of repetitive sequence structures in human centromeres. Science advances 2020, 6:eabd9230.

96. Fornes O, Castro-Mondragon JA, Khan A, Van der Lee R, Zhang X, Richmond PA, Modi BP, Correard S, Gheorghe M, Baranašić D: JASPAR 2020: update of the open-access database of transcription factor binding profiles. Nucleic acids research 2020, 48:D87–D92.

97. Grant CE, Bailey TL, Noble WS: FIMO: scanning for occurrences of a given motif. Bioinformatics 2011, 27:1017–1018.

98. Quinlan AR, Hall IM: BEDTools: a flexible suite of utilities for comparing genomic features. Bioinformatics 2010, 26:841–842.

99. Sha L, Yang Z, An S, Yang W, Kim S, Oh H, Xu J, Yin J, Wang H, Lenz HJ, et al: Non-canonical MLL1 activity regulates centromeric phase separation and genome stability. Nat Cell Biol 2023, 25:1637–1649.

