## Supplemental Data 1 for "Revealing long-range heterogeneous organization of nucleoproteins with N^6^-methyladenine footprinting"

Figure S1

A

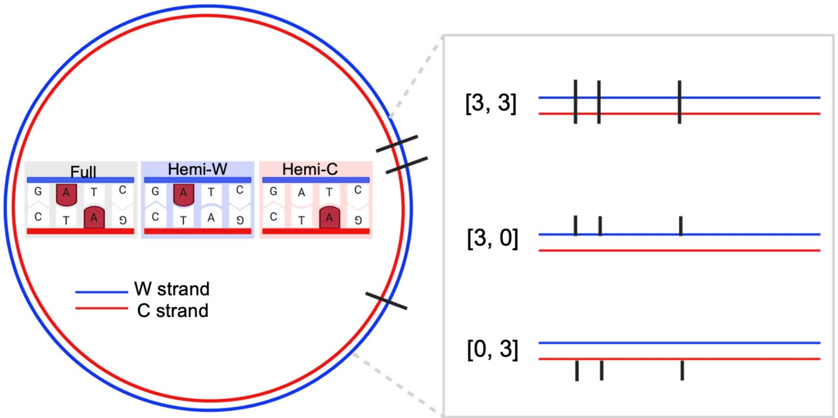


B

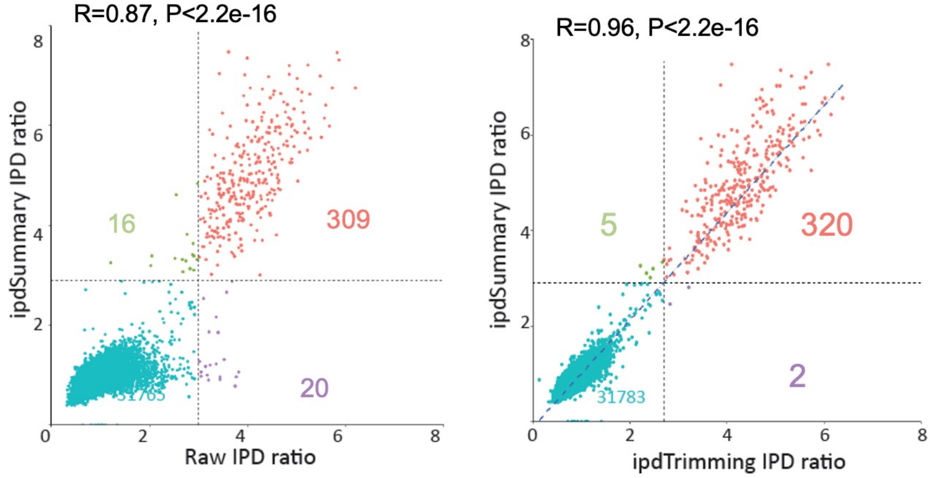


C

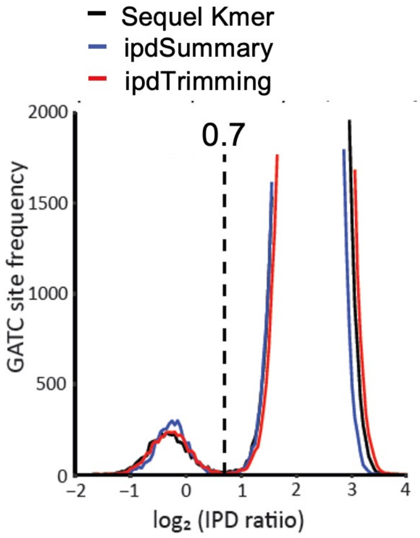


D

| Categories | Sequel Kmer | ipdSummary | ipdTrimming |
|---|---|---|---|
| [0, 1] | 2 | 2 | 2 |
| [1, 0] | 0 | 0 | 0 |
| [1, 1] | 0 | 0 | 0 |
| [0, 2] | 2 | 2 | 2 |
| [1, 2] | 2 | 3 | 2 |
| [2, 0] | 4 | 4 | 3 |
| [2, 1] | 0 | 0 | 1 |
| [2, 2] | 7 | 9 | 8 |
| [0, 3] | 178 | 183 | 182 |
| [1, 3] | 193 | 190 | 193 |
| [2, 3] | 542 | 538 | 535 |
| [3, 0] | 447 | 462 | 457 |
| [3, 1] | 310 | 307 | 309 |
| [3, 2] | 508 | 508 | 508 |
| [3, 3] | 50581 | 50568 | 50574 |

Figure S2

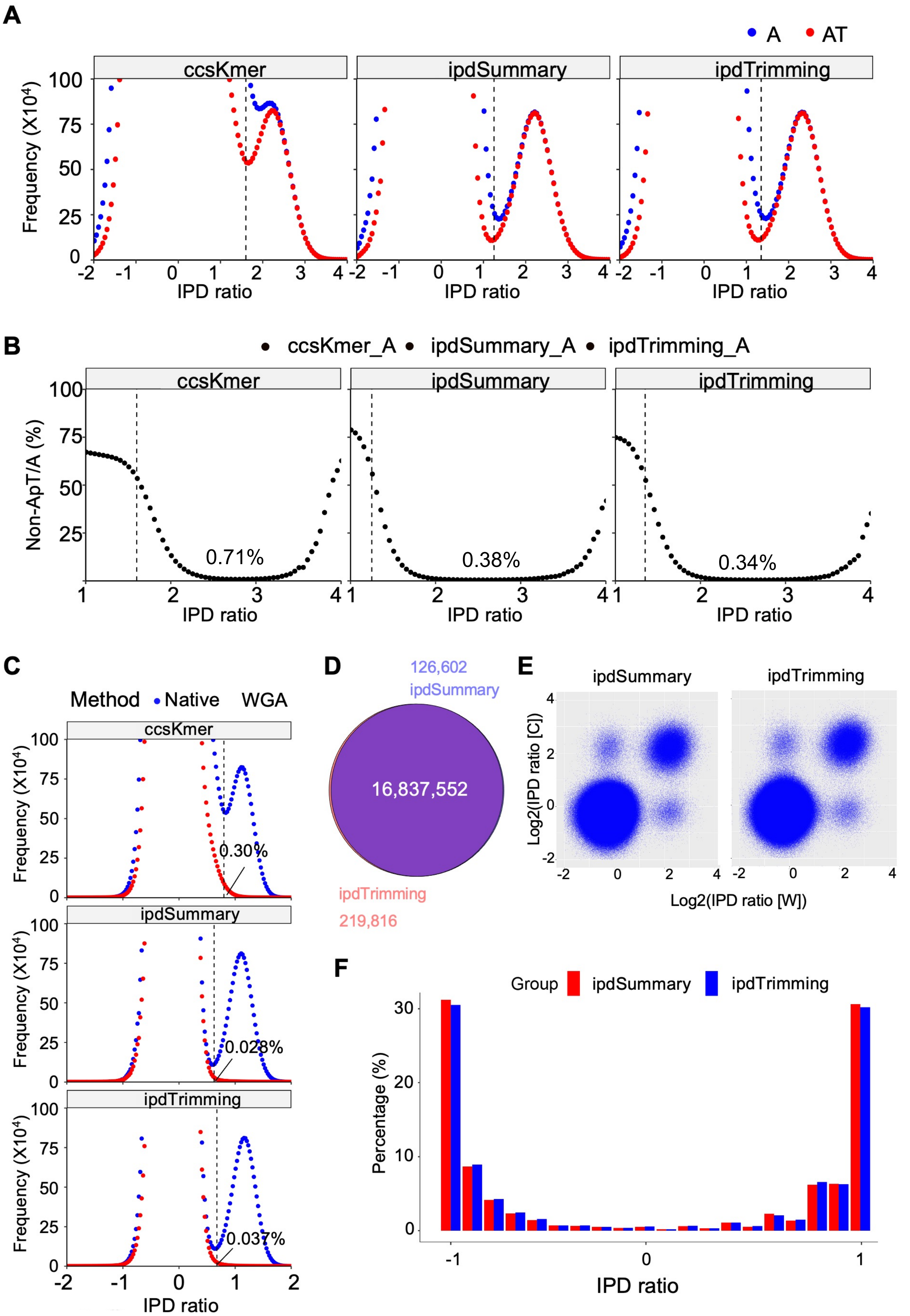
A
A
AT
ccsKmer
ipdSummary
ipdTrimming
Frequency (X10⁴)
IPD ratio
B
ccsKmer_A
ipdSummary_A
ipdTrimming_A
Non-ApT/A (%)
0.71%
0.38%
0.34%
IPD ratio
C
Method
Native
WGA
0.30%
0.028%
0.037%
D
126,602
ipdSummary
16,837,552
ipdTrimming
219,816
E
ipdSummary
ipdTrimming
Log2(IPD ratio [C])
Log2(IPD ratio [W])
F
Group
ipdSummary
ipdTrimming
Percentage (%)
IPD ratio

Figure S3

A

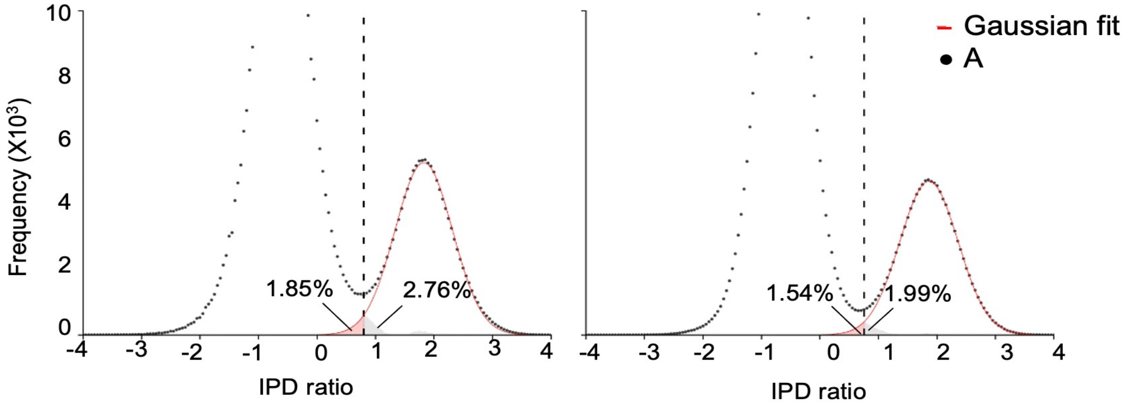


B

| Widom601 | Joint | | ipdSummary only | | ipdTrimming only | |
|---|---|---|---|---|---|---|
| | Count | Percentage(%) | Count | Percentage(%) | Count | Percentage(%) |
| linker | 638572 | 97.0 | 22722 | 85.9 | 13424 | 89.2 |
| nucleosome | 19615 | 3.0 | 3739 | 14.1 | 1619 | 10.8 |

C

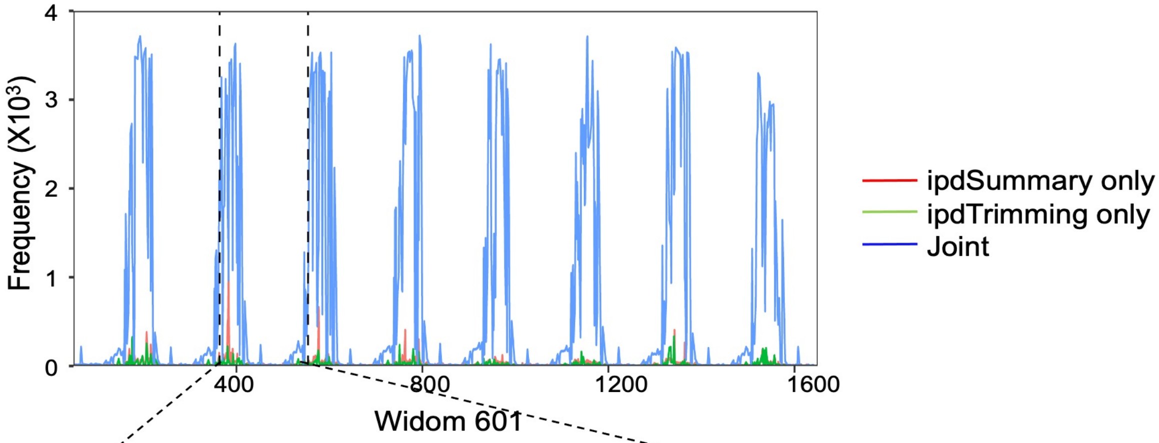


D

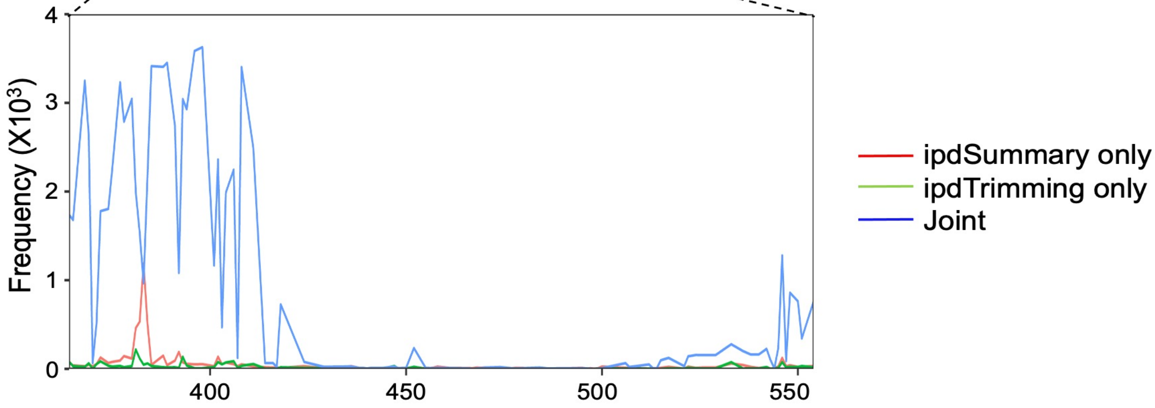

Figure S4

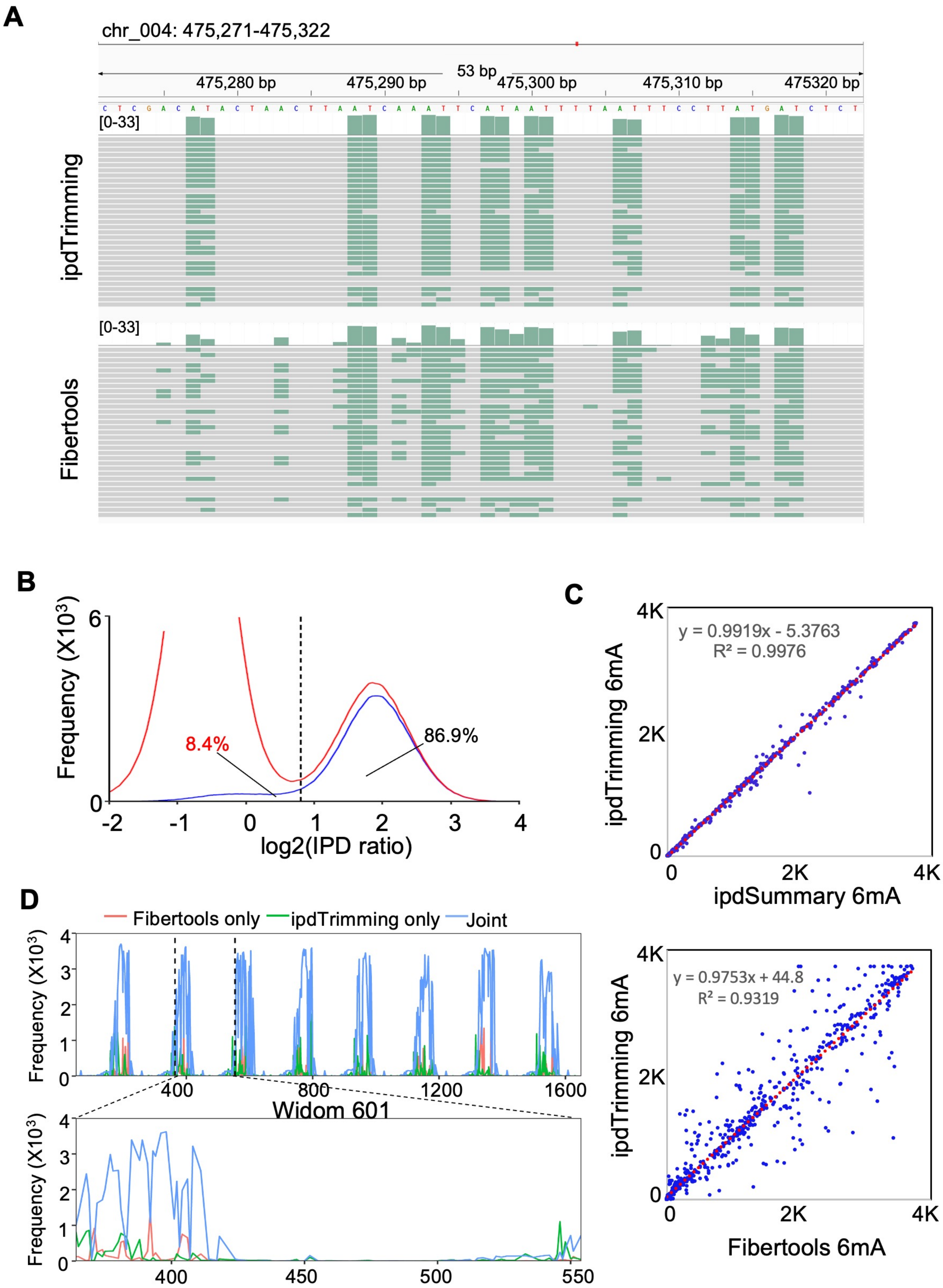
A
chr_004: 475,271-475,322
53 bp
475,280 bp
475,290 bp
475,300 bp
475,310 bp
475320 bp
[0-33]
ipdTrimming
[0-33]
Fibertools
B
Frequency (X10³)
8.4%
86.9%
log2(IPD ratio)
C
y = 0.9919x - 5.3763
R² = 0.9976
ipdTrimming 6mA
ipdSummary 6mA
y = 0.9753x + 44.8
R² = 0.9319
ipdTrimming 6mA
Fibertools 6mA
D
Fibertools only
ipdTrimming only
Joint
Frequency (X10³)
Widom 601

Figure S5

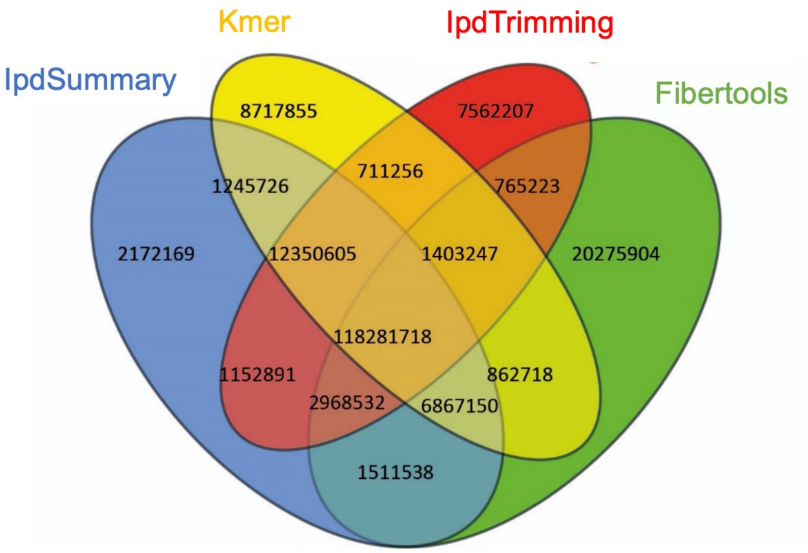

Kmer
IpdTrimming
IpdSummary
Fibertools
8717855
7562207
711256
1245726
765223
2172169
12350605
1403247
20275904
118281718
1152891
862718
2968532
6867150
1511538

Figure S6

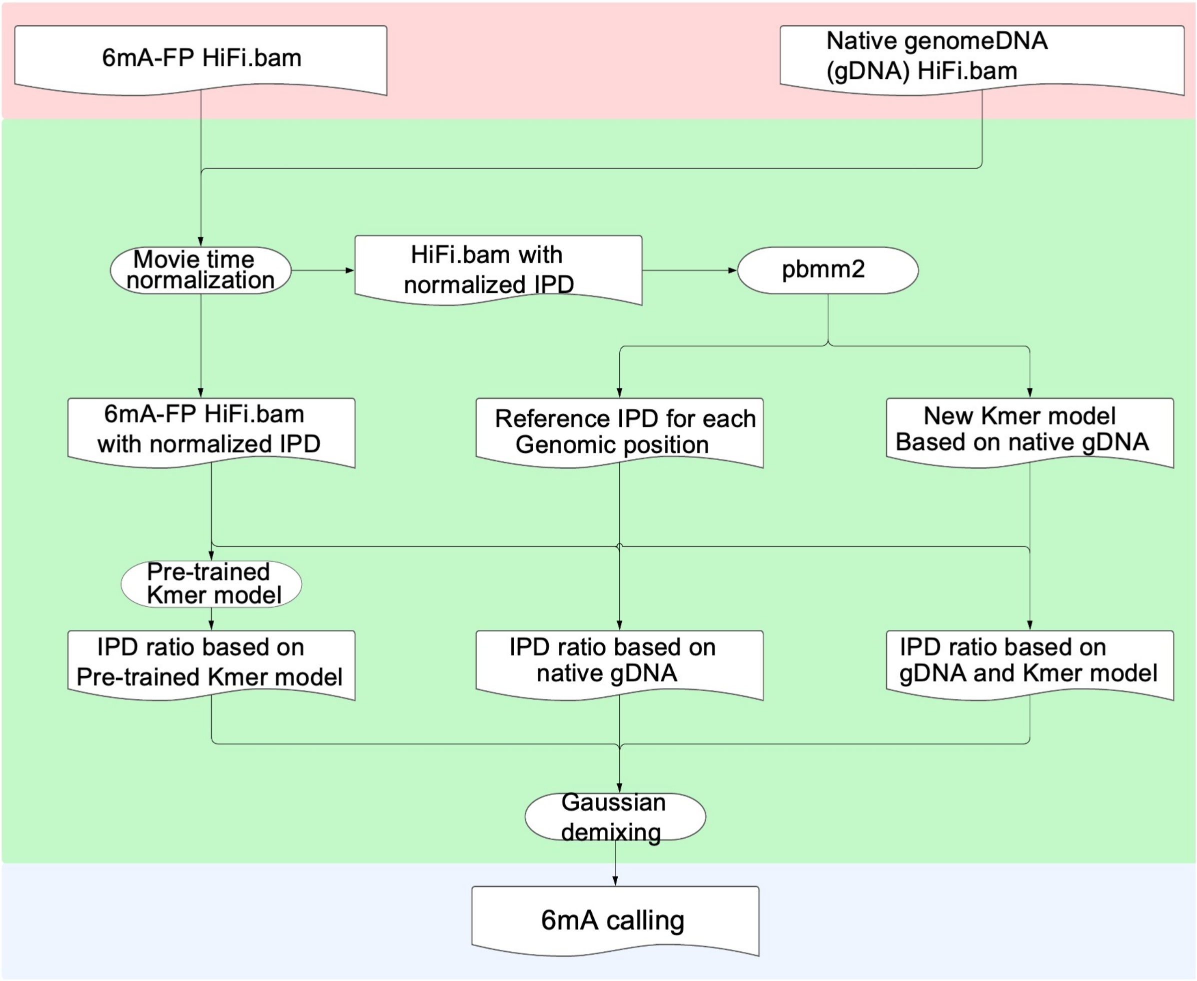
6mA-FP HiFi.bam
Native genomeDNA (gDNA) HiFi.bam
Movie time normalization
HiFi.bam with normalized IPD
pbmm2
6mA-FP HiFi.bam with normalized IPD
Reference IPD for each Genomic position
New Kmer model Based on native gDNA
Pre-trained Kmer model
IPD ratio based on Pre-trained Kmer model
IPD ratio based on native gDNA
IPD ratio based on gDNA and Kmer model
Gaussian demixing
6mA calling

Figure S7

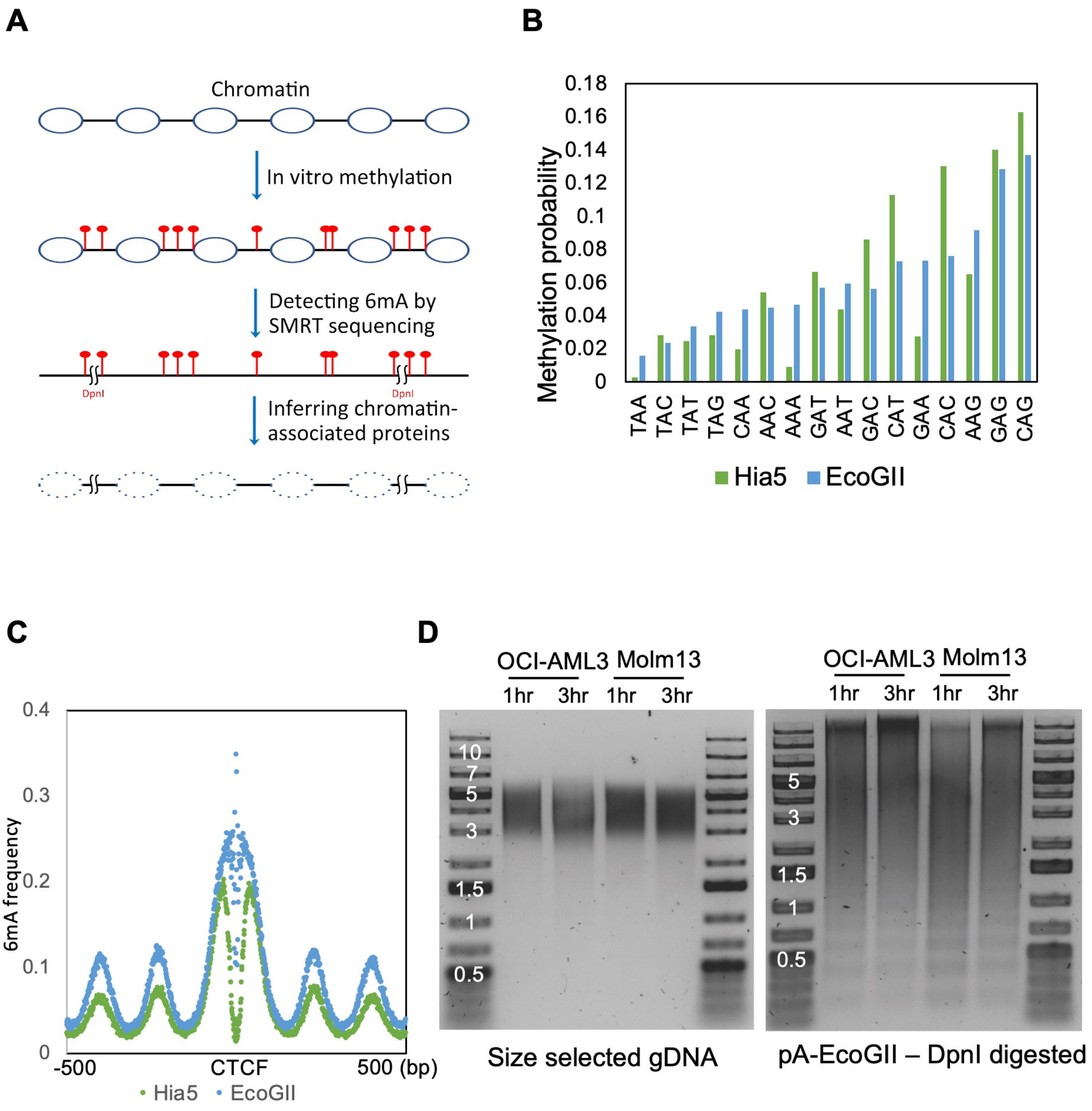
A
Chromatin
In vitro methylation
Detecting 6mA by SMRT sequencing
DpnI
DpnI
Inferring chromatin-associated proteins
B
Methylation probability
0.18
0.16
0.14
0.12
0.1
0.08
0.06
0.04
0.02
0
TAA TAC TAT TAG CAA AAC AAA GAT AAT GAC CAT GAA CAC AAG GAG CAG
Hia5
EcoGII
C
6mA frequency
0.4
0.3
0.2
0.1
0
-500
CTCF
500 (bp)
Hia5
EcoGII
D
OCI-AML3
Molm13
1hr
3hr
1hr
3hr
10
7
5
3
1.5
1
0.5
Size selected gDNA
OCI-AML3
Molm13
1hr
3hr
1hr
3hr
5
3
1.5
1
0.5
pA-EcoGII – DpnI digested

Figure S8

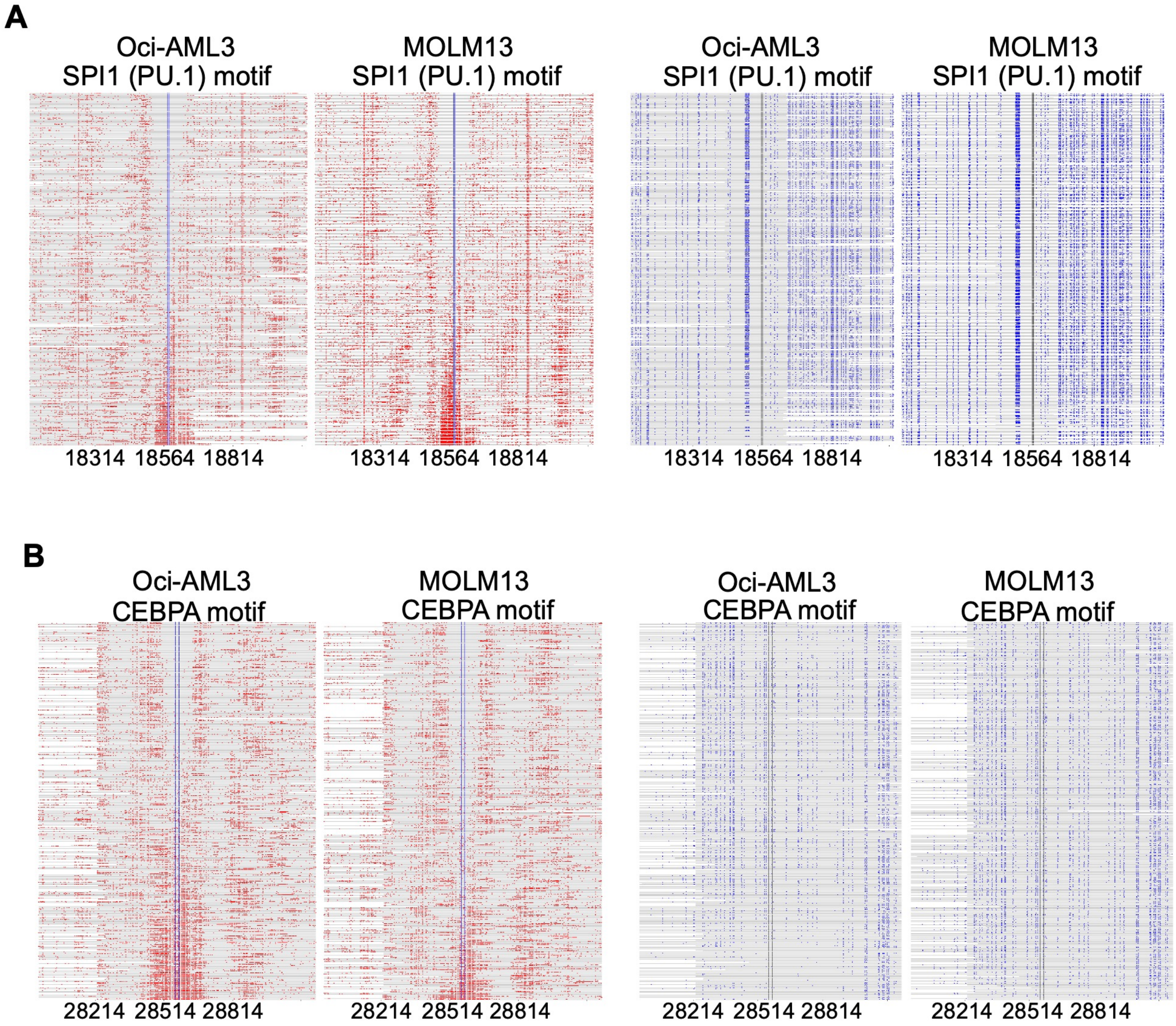
A
Oci-AML3
SPI1 (PU.1) motif
MOLM13
SPI1 (PU.1) motif
Oci-AML3
SPI1 (PU.1) motif
MOLM13
SPI1 (PU.1) motif
18314 18564 18814
18314 18564 18814
18314 18564 18814
18314 18564 18814
B
Oci-AML3
CEBPA motif
MOLM13
CEBPA motif
Oci-AML3
CEBPA motif
MOLM13
CEBPA motif
28214 28514 28814
28214 28514 28814
28214 28514 28814
28214 28514 28814

Figure S9

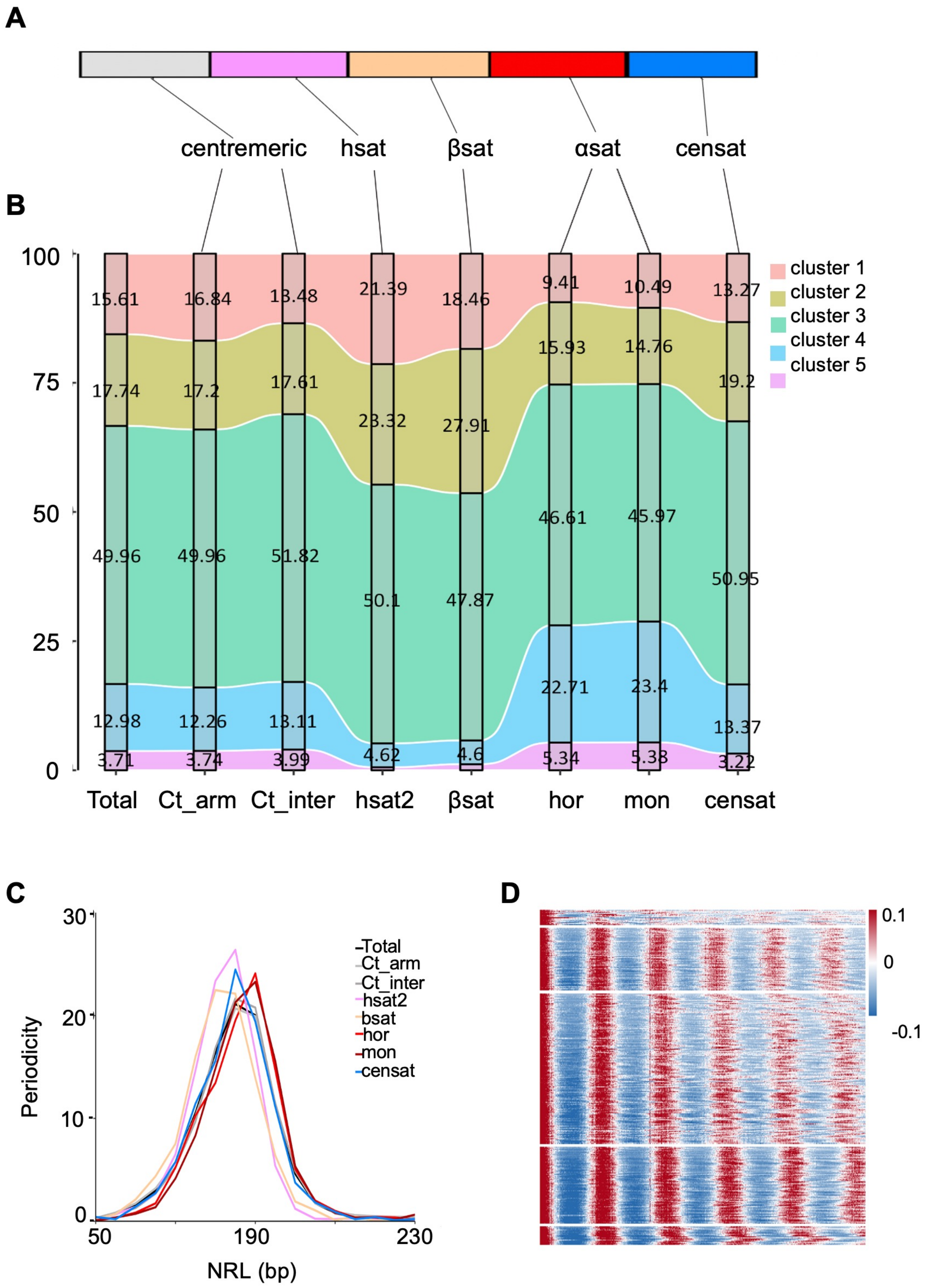
A
centremeric
hsat
βsat
αsat
censat
B
100
75
50
25
0
15.61
16.84
13.48
21.39
18.46
9.41
10.49
13.27
17.74
17.2
17.61
23.32
27.91
15.93
14.76
19.2
49.96
49.96
51.82
50.1
47.87
46.61
45.97
50.95
12.98
12.26
13.11
4.62
4.6
22.71
23.4
13.37
3.71
3.74
3.99
5.34
5.38
3.22
Total
Ct_arm
Ct_inter
hsat2
βsat
hor
mon
censat
cluster 1
cluster 2
cluster 3
cluster 4
cluster 5
C
Periodicity
30
20
10
0
50
190
230
NRL (bp)
Total
Ct_arm
Ct_inter
hsat2
bsat
hor
mon
censat
D
0.1
0
-0.1

Figure S10

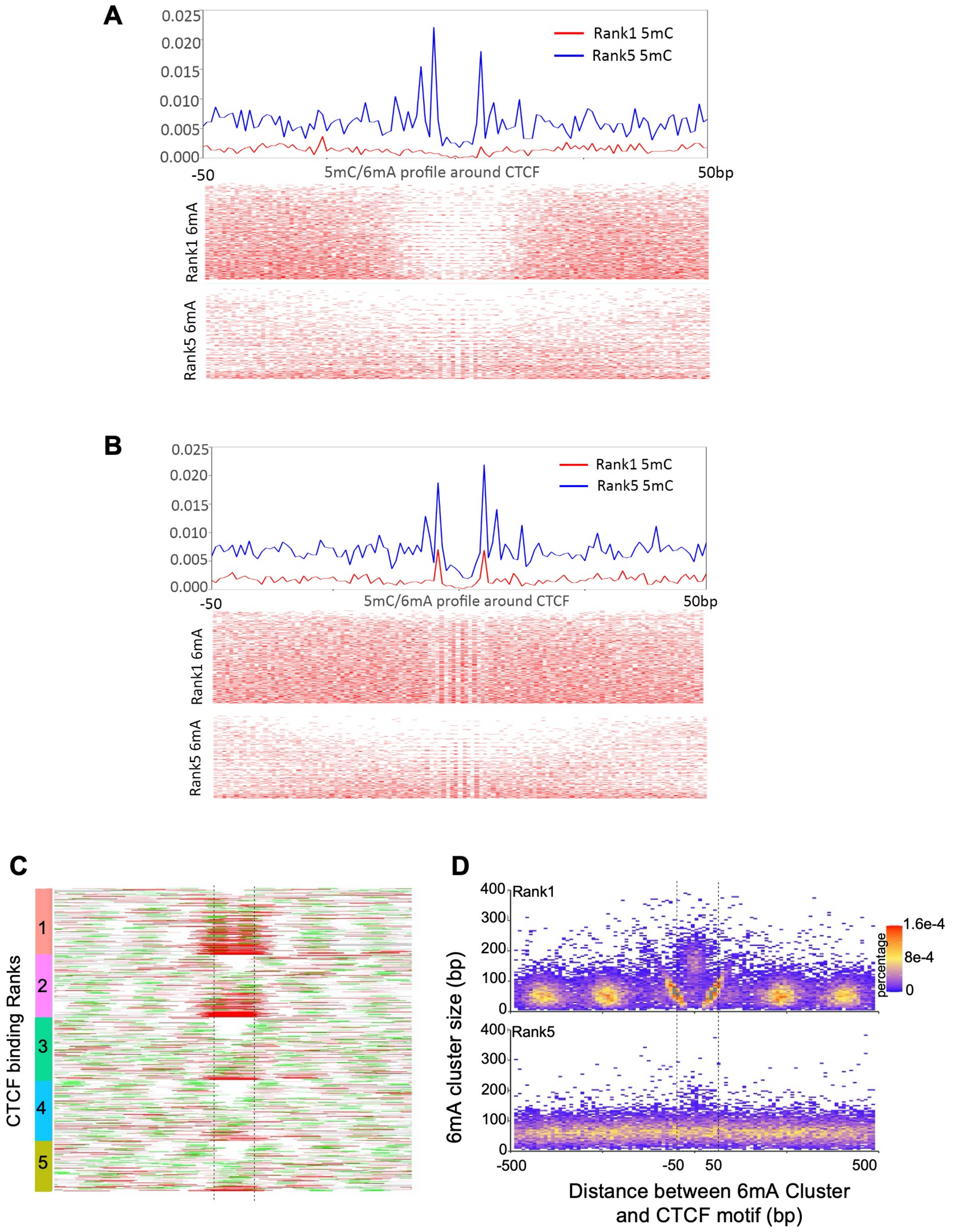
A
0.025
0.020
0.015
0.010
0.005
0.000
-50
50bp
5mC/6mA profile around CTCF
Rank1 5mC
Rank5 5mC
Rank1 6mA
Rank5 6mA
B
0.025
0.020
0.015
0.010
0.005
0.000
-50
50bp
5mC/6mA profile around CTCF
Rank1 5mC
Rank5 5mC
Rank1 6mA
Rank5 6mA
C
CTCF binding Ranks
1
2
3
4
5
D
Rank1
Rank5
400
300
200
100
0
6mA cluster size (bp)
1.6e-4
8e-4
0
percentage
-500
-50
50
500
Distance between 6mA Cluster and CTCF motif (bp)

Figure S11

A

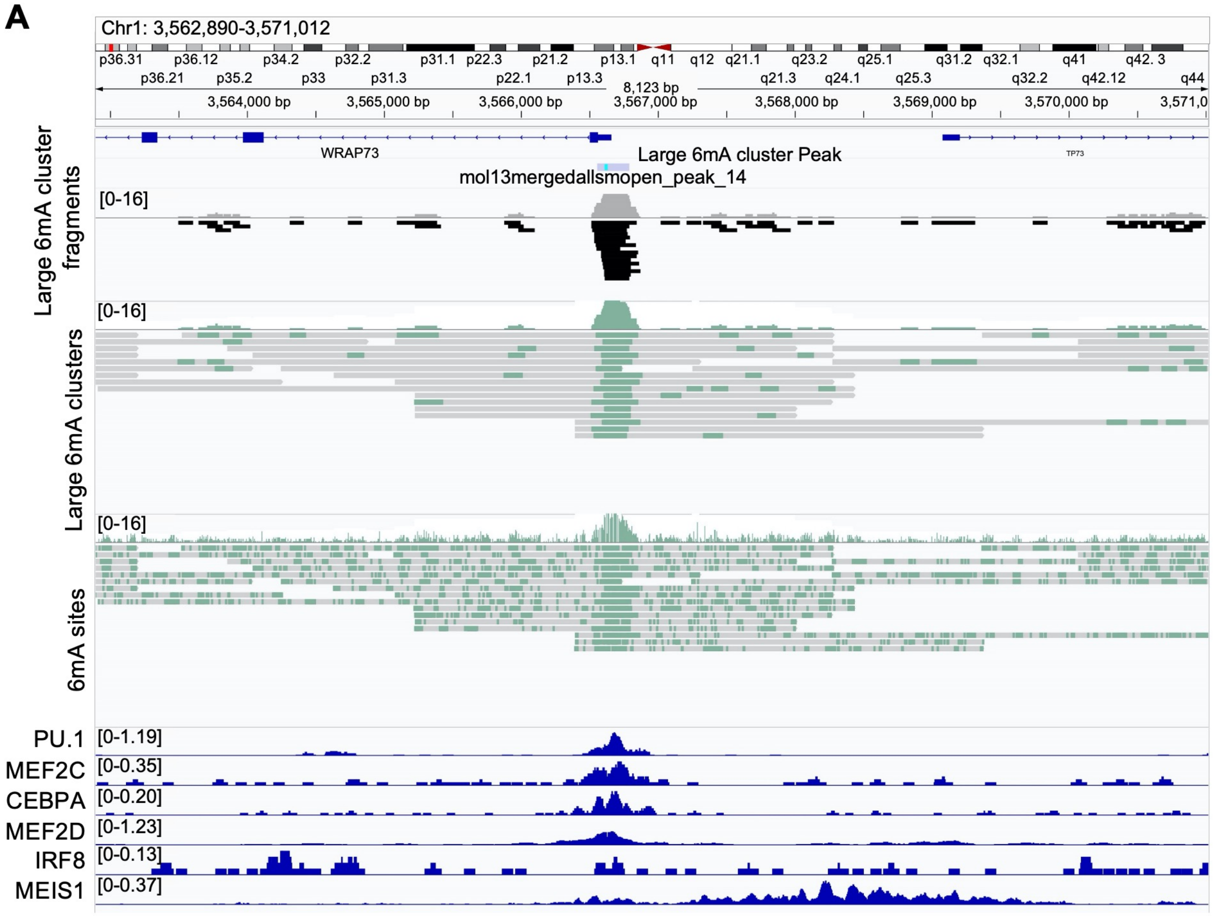


B

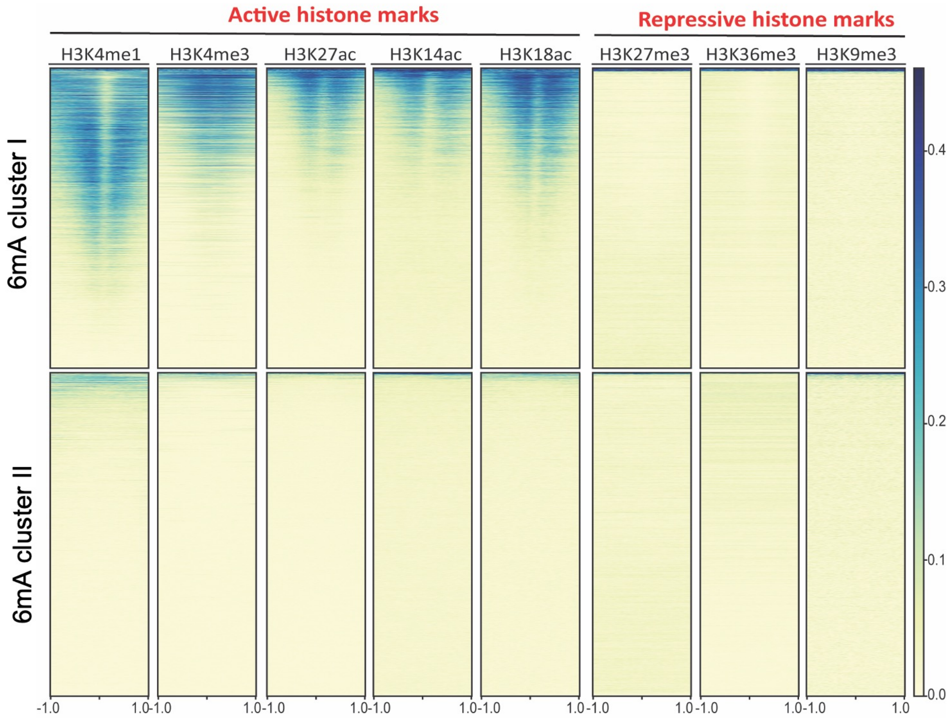

Figure S12

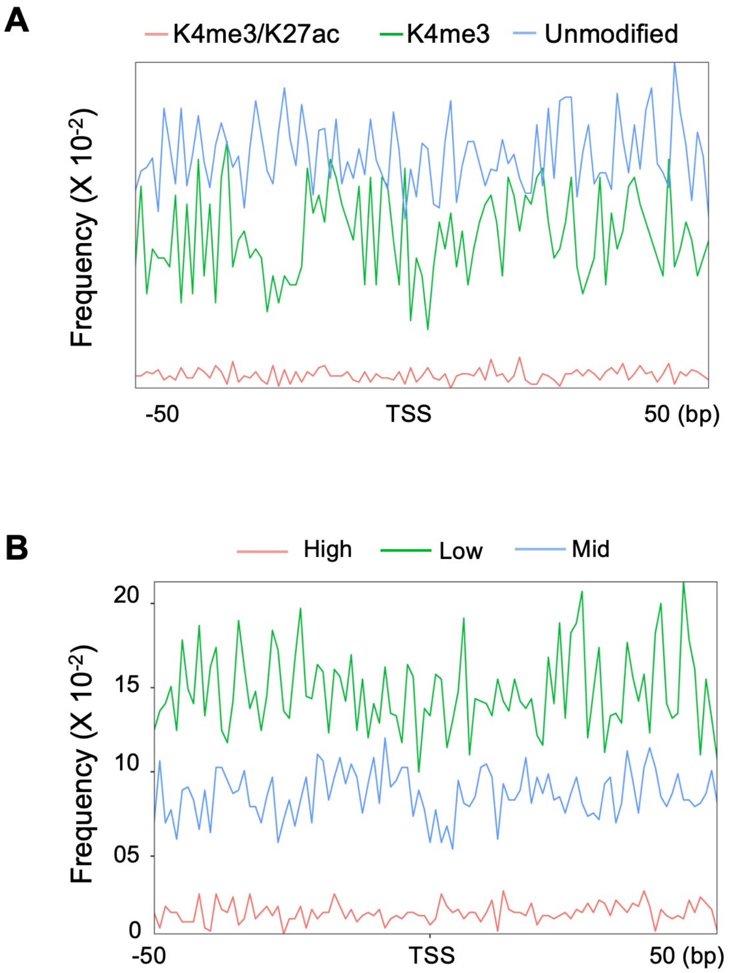
A
K4me3/K27ac
K4me3
Unmodified
Frequency (X 10-2)
-50
TSS
50 (bp)
B
High
Low
Mid
Frequency (X 10-2)
20
15
10
05
0
-50
TSS
50 (bp)

Figure S13

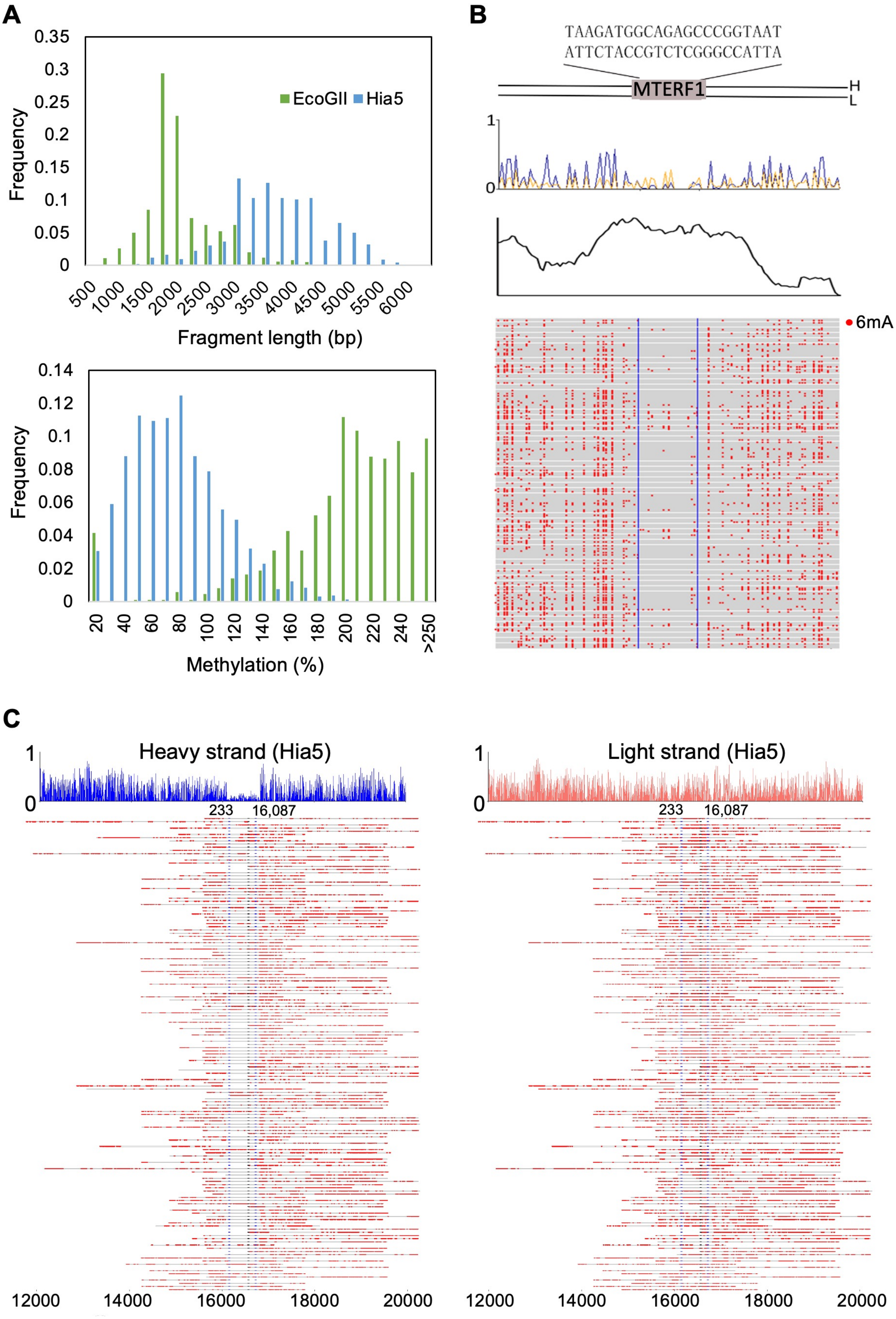

A
EcoGII
Hia5
Frequency
Fragment length (bp)
Methylation (%)
B
TAAGATGGCAGAGCCCGGTAAT
ATTCTACCGTCTCGGGCCATTA
MTERF1
H
L
6mA
C
Heavy strand (Hia5)
Light strand (Hia5)
233
16,087
